# Hippocampal gamma and sharp wave/ripples mediate bidirectional interactions with cortical networks during sleep

**DOI:** 10.1101/2022.03.08.483425

**Authors:** Rafael Pedrosa, Mojtaba Nazari, Majid H. Mohajerani, Thomas Knöpfel, Federico Stella, Francesco Battaglia

**Author notes:** **Correspondence:** Francesco Battaglia, Ph.D., Federico Stella, Ph.D., Rafael Pedrosa M.Sc, Radboud University, Donders Institute for Brain, Cognition and Behaviour, Neuroinformatica, Heyendaalseweg 135, 6525 AJ NIJMEGEN, Internal postal code: 66. Equal contribution (Federico Stella, Francesco Battaglia).

## Abstract

Hippocampus-neocortex interactions during sleep are critical for memory processes: hippocampally-initiated replay contributes to memory consolidation in the neocortex and hippocampal sharp wave/ripples are linked to generalized increases in neocortical cell activity and DOWN-UP state transitions. Yet, the spatial and temporal patterns of this exchange are unknown. With voltage imaging, electrocorticography, and laminarly-resolved hippocampal potentials, we characterized cortico-hippocampal interactions during anesthesia and NREM sleep. We observed neocortical activation transients spanning multiple spatial scales hinting at a quasi-critical regime. Transients were organized in a small number of functional networks matching known anatomical connectivity. A network overlapping with the default mode network and centered on retrosplenial cortex was the most associated with the hippocampus. Interestingly, hippocampal slow gamma was the oscillation that best correlated with this neocortical network, outpacing ripples. In fact, neocortical activity predicted hippocampal slow gamma and followed ripples, suggesting that consolidation processes rely on bi-directional exchanges between hippocampus and neocortex.

## Introduction

Spontaneous activity during quiescent periods and sleep is likely to be crucial for a host of cognitive functions such as imagery, planning, self-reflection (Andrews-Hanna, 2012; Raichle, 2015). As supported by evidence from interventional studies (Genzel et al., 2014; Girardeau et al., 2009; Maingret et al., 2016; Marshall et al., 2006) offline activity plays a decisive role in memory consolidation, the strengthening, stabilization and re-organization of memories for the long term (Dudai, 2004; Frankland and Bontempi, 2005). For this, the dialogue between the hippocampus and the neocortex seems to be relevant. These two brain structures were postulated to form two complementary memory systems (McClelland et al., 1995), with the hippocampus forming rapid memories and “index codes” that summarize and point to brain-wide activity patterns (Teyler and DiScenna, 1986). In this scheme, the neocortex would be the larger storage containing information forming a statistical model of the world, integrating the totality of the remembered experience(Battaglia et al., 2012; McClelland and Rogers, 2003).

Memory replay, initiated by the hippocampus, is considered as a potential conduit for the transmission of the “index code”. Replay is the spontaneous repetition of neural activity patterns that were initially elicited during experience (Battaglia et al., 2011; O’Neill et al., 2010). In wakefulness, replay may contribute to abilities like planning and mental simulation (Kaefer et al., 2020; McNamara et al., 2014; Pfeiffer and Foster, 2013). During sleep, it has been hypothesized to enable the merging of hippocampal memories into the cortical store, gradually enough to prevent “catastrophic interference” and the loss of memory(McClelland et al., 1995). Hippocampal sharp waves/ripples (SWR) (Buzsaki, 2015), a burst of activity which coincides with the bulk of replay in the hippocampus, have been posited to act as the carrier of that information. During NREM sleep, cortical activity is dominated by large transient fluctuations in activity, the so-called UP-DOWN states (Steriade et al., 1993a). The elevated activity component of the fluctuation, the “UP” state, is likely maintained by recurrent excitation in the local cortical network neurons (Shu et al., 2006; Steriade et al., 1993b). Importantly, hippocampal SWRs correlate with the DOWN to UP state transition (Battaglia et al., 2004), an interrelationship that may be key to memory consolidation, because SWRs correlated with increased cortical replay (Ji and Wilson, 2007; Peyrache et al., 2009) and DOWN-UP state transitions temporally organize replay (Johnson et al., 2010; Peyrache et al., 2009).

While the association between SWRs, hippocampal replay and replay in a handful of cortical areas tightly connected with the hippocampus, like medial prefrontal cortex (Peyrache et al., 2009, 2011), retrosplenial (Nitzan et al., 2020), entorhinal (Olafsdottir et al., 2016) has been well documented, how hippocampal and cortex-wide activity relate is less known. Generalized increases in cortical activation during hippocampal SWRs were found with single cell electrophysiology (Battaglia et al., 2004), functional MRI (Logothetis et al., 2012), and voltage and glutamate wide-field Imaging (Karimi Abadchi et al., 2020), however, a detailed account of the dynamics of the interactions between large cortical networks and the hippocampus is still missing and this is what we set out to provide here. This is especially important since systems consolidation (McClelland et al., 1995) and index theory (Teyler and DiScenna, 1986) assume whole-cortex activity configurations as the substrate of memories.

UP/DOWN state fluctuations are coherent across large swaths of cortex, and have often been described as traveling waves (Amzica and Steriade, 1995a; Huber et al., 2004; Mohajerani et al., 2010, 2013). A complementary description emphasizes that transient activations may span a very wide range of scales, from local to whole cortex (Mohajerani et al., 2010). Voltage sensitive imaging from an offline state (recovery from anesthesia) (Scott et al., 2014) showed a size distribution for transients that approximates a power-law, the signature of criticality. Criticality (or near-criticality (Wilting and Priesemann, 2019)) is a dynamical state that maximizes long-range correlations, and has potential advantages for computational power, and can be precisely defined and quantified by a number of estimators (Wilting and Priesemann, 2018). Yet, a fully statistical discussion that ignores the highly structured nature of cortical connectivity would not be very useful in unraveling the dynamical basis of memory. Here, we used voltage imaging and electrocorticography in mice to provide a wide-scale picture of spontaneous cortical activity in offline states, and decomposed them in functional networks. Closely matching the results of a large anatomical projections dataset (Zingg et al., 2014), our data-driven procedure identified three networks, centered respectively on the retrosplenial cortex and medial cortical bank of the cortex, on somatosensory cortex and on lateral cortex.

We then set out to analyze the hippocampal CA1 layer-resolved LFP correlates of transients involving these three cortical networks. We did not restrict our analysis to SWR events, but considered hippocampal oscillations in other frequency bands as well. Strikingly, Slow Gamma (20-50 Hz) centered in CA1 stratum radiatum was as strongly, if not more, correlated with cortical activity than SWR, and with the retrosplenial network in particular. Slow Gamma is thought to be related to the routing of information from CA3, a recurrent-rich hippocampal subfield, which for this reason is seen as potentially functioning as an auto-associative memory, and CA1 (Bragin et al., 1995; Colgin et al., 2009). Consistent with this idea, Slow Gamma in wakefulness is related in CA1 with an increase in the predictive features of spatially-related cell activity (Bieri et al., 2014; Cabral et al., 2014; Lasztóczi and Klausberger, 2016) and with the organization of neuronal spiking into ordered sequences (Guardamagna et al., 2021). During sleep, increased Slow Gamma during SWR events correlates with greater replay (Carr et al., 2012). Together, this evidence points at a role for Slow Gamma in memory recollection in CA1. Our results suggest that spontaneous activity in the cortex may act as a “cue” for such retrieval, and characterize the cortico-hippocampal exchange as a bi-directional dialogue. Such interaction may be necessary for creating consistent index representations (Mcnaughton et al., 2002; Teyler and DiScenna, 1986) and providing the basis for extending complementary learning systems theory to the entirety of the neocortex. Interestingly, the retrosplenial network, that is the most involved in this exchange with the hippocampus, strongly overlaps with the Default Mode Network (Kaplan et al., 2016; Stafford et al., 2014). Thus, our data also provides a potential dynamical scenario explaining the involvement of the DMN in memory.

## Results

### Parallel recordings of neural activity transients in the cortex and the hippocampus

We imaged the neocortical spatiotemporal activity in the right hemisphere (Barson et al., 2020; Mohajerani et al., 2010; Ren and Komiyama, 2021; Song et al., 2018) under urethane anesthesia and natural sleep (Figure 1 A–B, Figure S1). For this, we used wide-field macroscopy and a voltage-sensitive dye (VSD) in wild-type C57/Bl6 mice (Bermudez-Contreras et al., 2018; Karimi Abadchi et al., 2020; Mohajerani et al., 2013) (for the anesthesia experiments) and mice expressing a ratiometric genetically encoded voltage indicator (GEVI; chiVSFP (Akemann et al., 2010, 2012; Carandini et al., 2015; Song et al., 2018); for natural sleep, here referred to as VSI). In both cases, the neocortical voltage activity was simultaneously recorded with the ipsilateral hippocampal Local Field Potential (LFP; an electrode was placed in dorsal CA1 for the urethane anesthesia recordings and a 16 channels high density silicon probe recorded signals from all CA1 layers for natural sleep recordings). In the urethane anesthesia experiments, we also sampled cortical electrical activity by use of an electrocorticography (ECoG) 6×5 grid placed on the same cortical surface we were imaging. In neocortex, we observed temporal activity fluctuations spanning a large range of temporal and spatial scales (Figure 1 C). We then applied an activity transient detection algorithm based on local and global average neocortical activity (Figure 1 D-E, see the section “Cortical transient detection” on methods). Such method allowed us to isolate temporal windows of excitation spanning variable neocortical extents and to separately analyze the global properties of these activation bouts, such as their duration and size.

**Figure 1:**
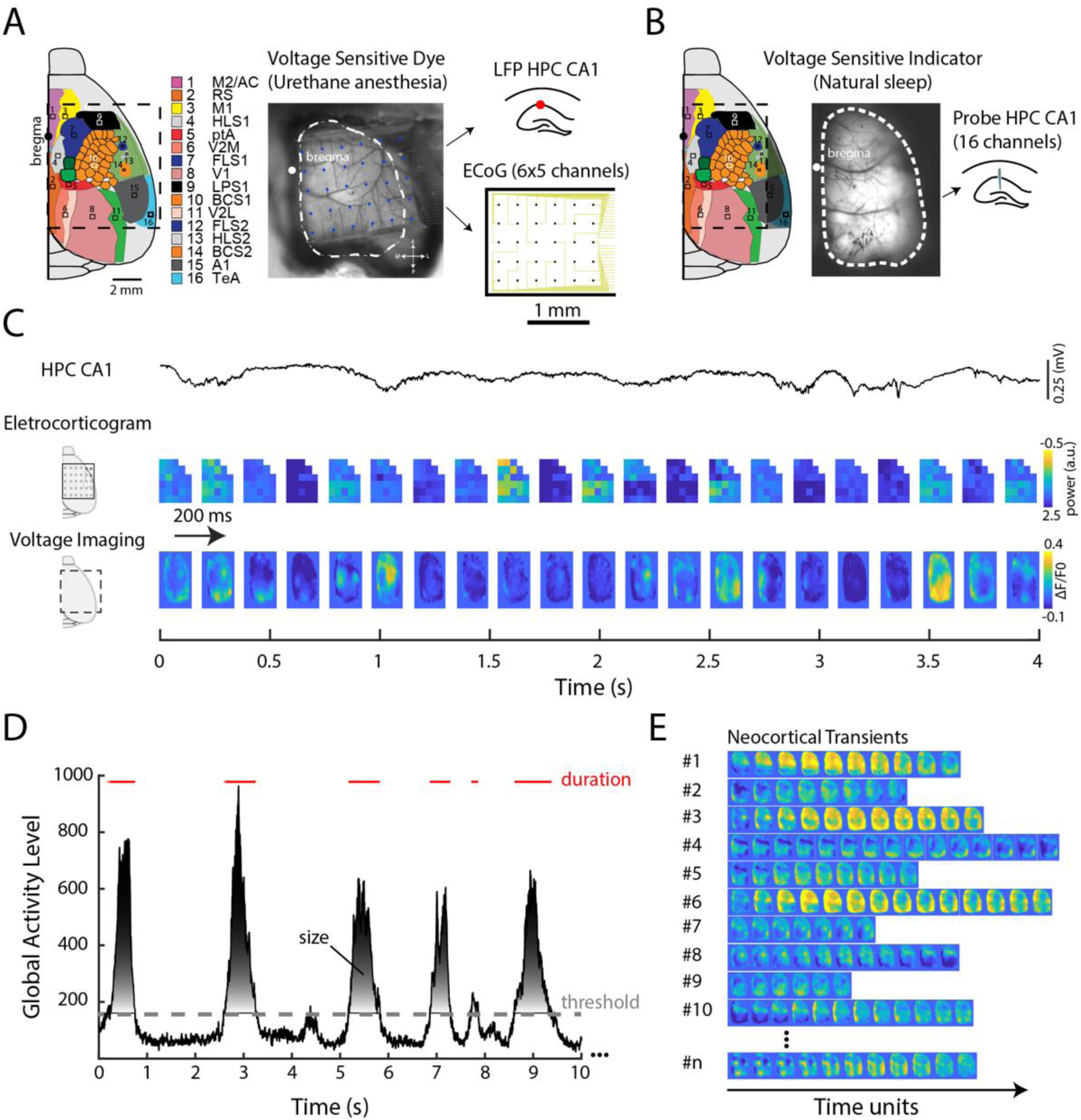
Experimental setups used for the neocortical transients recording. **(A)** Schematic of wide-field topography (left) and photo of the preparation (center) of the setup for the neocortical voltage sensitive dye imaging recorded in combination with hippocampal LFP and ECoG in mice under urethane anesthesia. The black dashed line in the diagram (white dashed line in the photo) represents the borders of the camera field of view. Bregma is represented as a black dot. Insets to the right depict the transparent ECoG (6×5 channels) covering much of the imaging field of view, and the location of hippocampal LFP electrode. **(B)** Schematic and photo of ipsilateral wide-field topography of the neocortical voltage sensitive indicator imaging recorded in combination with a linear silicon probe (16 channels) in the hippocampus. In this experiment, we recorded natural sleep in genetically encoded voltage indicator (GEVI) mice. Note that imaged region is smaller than in the anesthesia group (because skull was not removed, but only thinned). **(C)** Example traces of hippocampal LFP, ECoG, and VSD signals (anesthesia group). **(D)** Global activity level of the neocortex recorded by the VSD. The shadowed area represents the size of the transient, defined by the area under the curve, and red lines highlight the intervals relative to each transient. **(E)** Examples of detected neocortical transients.

### Cortical transients in the mouse cortex show power-law-like distributions of size in anesthesia and natural sleep

To characterize the properties of neocortical activity transients, we computed the statistical distribution of transient descriptors such as the overall, cumulative activation of cortex over the duration of the transient (Figure 2 A, Figure S2). In both urethane anesthesia and natural sleep groups we find that the frequency distribution of magnitude-related properties (size, duration) of transients are well captured by a power-law. We nevertheless observe a substantial difference in the goodness-of-fit between urethane anesthesia and natural sleep recordings: power-law fits were generally better for sleep data, mostly because anesthesia was associated with a pronounced over-abundance of large-sized events (and similarly of longer events), making the overall distribution resemble a supercritical one (Scott et al., 2014) (Urethane anesthesia/VSD group: α = −1.8775 and ε = 0.2325 and natural sleep/VSI group: α = −1.7385 and ε = 0.0856). Interestingly, such supercriticality is not observed in ECoG recording of the same cortical states (Urethane anesthesia/ECoG group: α = −2.168 and ε = 0.1751 and Urethane anesthesia/ECoG in VSD group: α = −1.8456 and ε = 0.2106), suggesting a substantial difference in the signal represented in the two recordings methods. Based on theoretical arguments (Neto et al., 2020; Nonnenmacher et al., 2017; Touboul and Destexhe, 2017) this may be due to the non-independence of electrode in the ECoG matrix, because of volume conductance effects. Although different in their statistical properties, VSD and ECoG signals were still strongly correlated, consistently with them being two measures of the same underlying phenomenon (Figure 2 B, Figure S2 B-D; Size correlations: mouse#1 = 0.65, mouse#2 = 0.61, mouse#3 = 0.54, mouse#4 = 0.42 and mouse#5 = 0.5).

**Figure 2:**
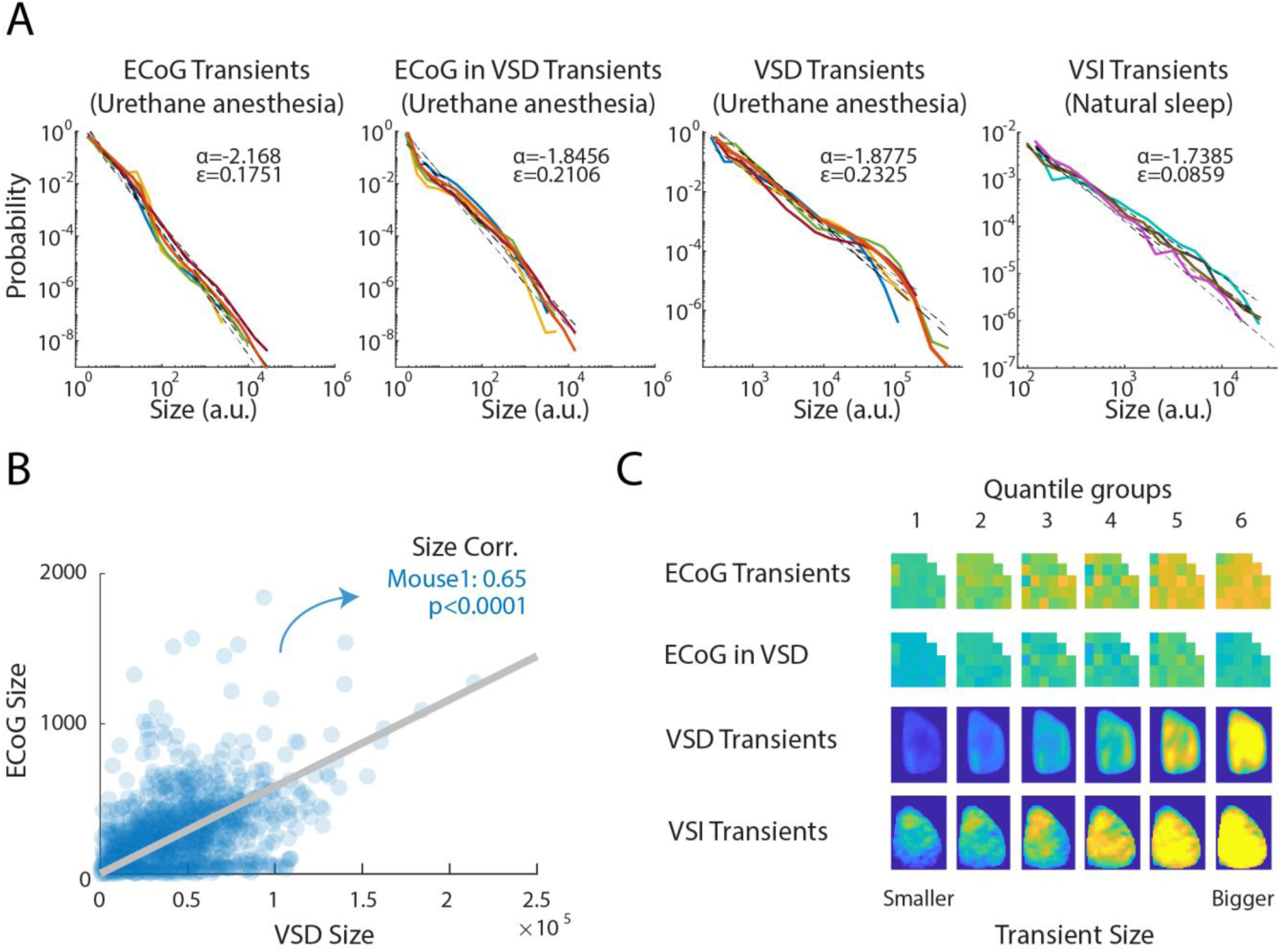
Probability distribution of transient sizes in urethane anesthesia and natural sleep (in log-log scale, a straight line corresponds to a power-law distribution). **(A)** Transient probability size in urethane anesthetized and natural sleep states. For the urethane dataset, we computed the size of ECoG transients detected by the negative deflection of ECoG data (i), size of ECoG transient associated to events detected from VSD data (ii) and the size of transients detected and measured from the VSD data (iii). For the natural sleep dataset, we computed neocortical transients of the VSI data during the NREM periods (iv, see Materials and Methods for the sleep state detection). Lines of different colors represent different animals. Note that the animals from the natural sleep code have a different color code. The black dashed lines represent the linear regression (in log-log scale) for each animal. α is the critical exponent (slope of the regression line) and ε the fitting error (see Methods) **(B)** Scatter plot of the transient sizes detected by the VSD data and the size of ECoG transients taking place at the same time, for mouse 1. **(C)** Neocortical average signals for the transients subdivided in 6 quantiles based on the size shown in panel (A).

### A few cortical networks account for small cortical transients

While size distributions provide information on the relative abundance of smaller vs. larger, global transients, they are not informative about the spatial organization of their events, or on whether different type of transients exist that span different cortical regions.

To address these questions, we first segmented the neocortical transients in 6 equal quantiles based on their size (Figure 2C). Then we classified the spatial patterns of activity by training a Restricted Boltzmann Machine (RBM) on temporal frames belonging to each of these transient quantiles. The RBM encodes these images based on the statistics of pixels co-activations and the state of each hidden unit state becomes selective for different a recurring arrangement of active pixels. To ascertain whether transient events can be classified in a finite number of discrete categories, we applied a clustering algorithm on the set of learned RBM weights (Köster et al., 2014) relative to all events (Figure 3 A, Figure S3 F). We find a marked dependency on the transient size. In both urethane anesthesia and natural sleep groups, transients at the lowest end of the quantile distribution can in fact be neatly subdivided in a very low number of well separated modes (scatter plots in Figure 3A; EM model fitting resulted in three modes as being the most likely subdivision for 4 out of 5 animals for the urethane anesthesia group and 4 out of 5 animals for the natural sleep group. The cluster separation was confirmed by the strong bimodal distribution of pairwise distances between RBM weight vectors as shown in the histograms). For larger transients, no clearly distinct modes appeared.

**Figure 3:**
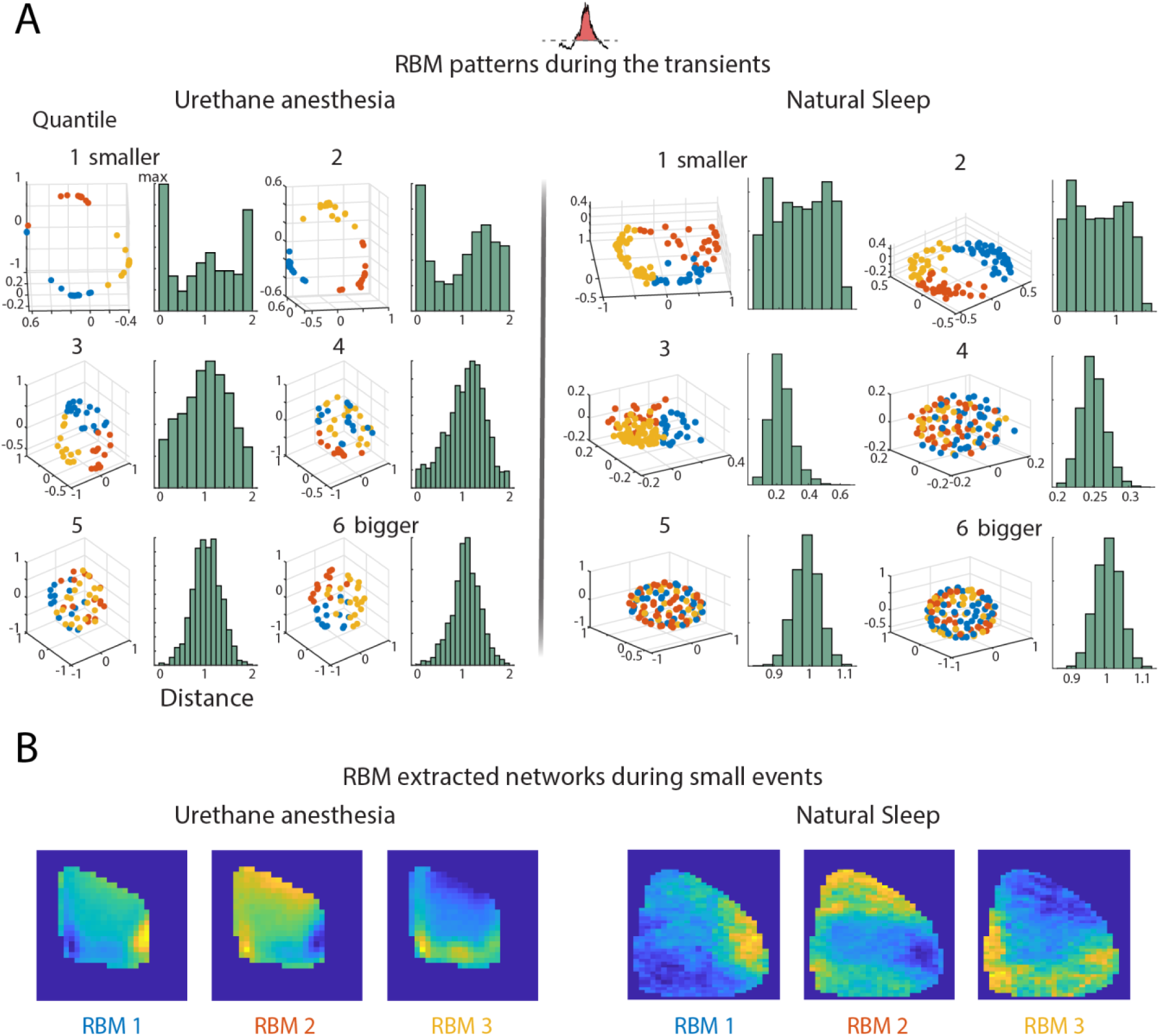
RBM reveals modular networks specifically for small neocortical transients. **(A)** Example of RBM weights (reduced to three dimensions by PCA) computed on each of the 6 quantile groups distributed by the transient size for urethane anesthesia and natural sleep data. The blue, orange and yellow colors denote 3 clusters separated by using the k-means algorithm. The respective green histograms show the distribution of cosine distances between hidden unit states for individual transients. Note that for both datasets a tendency for clustering in the RBM weights can be observed in smaller transient groups. **(B)** Spatial distribution of activity corresponding to the 3 k-means clusters in the first quartile RBM weights.

While large activations consist of the simultaneous activation of large swaths of cortical surface, small ones disproportionally involve a handful of networks centered around, respectively: Retrosplenial Cortex (RS), Temporal association area (TeA) and Somato-Sensory (SS) (Figure 3B). Further inspection of the spatial organization of the RBM-identified modes, reveals a tendency for these areas to form pairwise activation patterns (Figure S3 G). Thus, a set of well-defined networks appear to dominate the small activity fluctuations which make for the vast majority of transient cortical activity.

### Large cortical transients are most often preceded by RS or SS activation

As seen from the analysis above, large neocortical transients in both urethane anesthesia and natural sleep could not be straightforwardly classified in a handful of modes. In fact, activity propagation during these periods do not seem to follow stereotypical dynamical patterns (Video 1). Instead, we actually found that large transients (in particular those in the sixth and largest quantile) mostly differ by the activation patterns immediately preceding them. When applying the same previous RBM-based analysis on time frames extracted from short periods (∼100ms) immediately preceding the detected onset of the large transient, we found activation modes resembling those of Figure 3, with activations preceding last-quantile transients neatly clustering into a couple of well separated groups (EM model fitting resulted in two modes as being the most likely subdivision for: 5 out of 5 animals for the urethane anesthesia group and 3 out of 5 animals for the natural sleep group; Figure 4A). This is however not the case for smaller transients.

**Figure 4:**
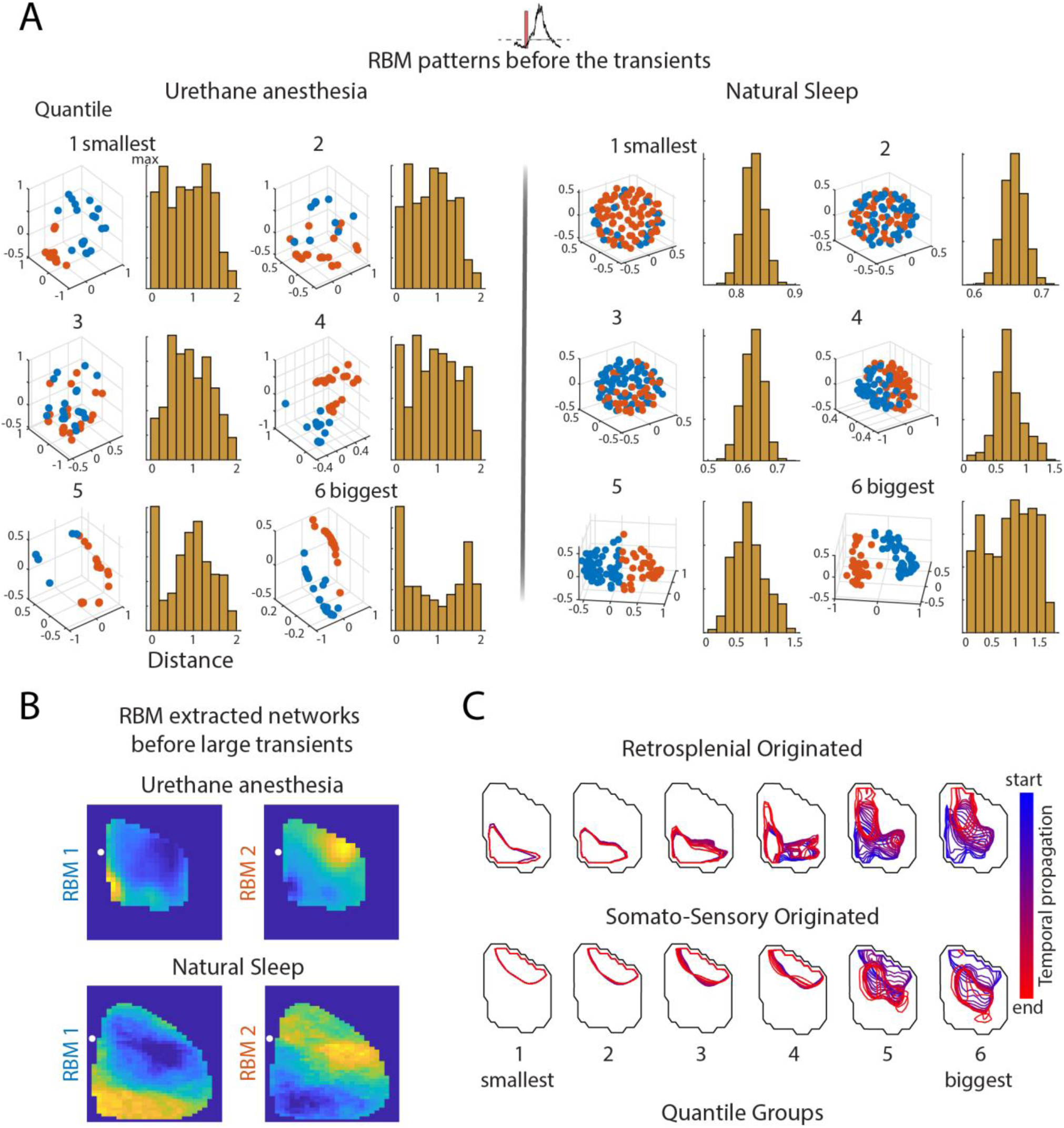
Large transients are triggered by Retrosplenial and Somato-Sensory cortex activation. **(A)** Example of RBM weights computed from cortical activations 100 ms before the transient onset for each of the 6 size quantiles for urethane anesthesia and natural sleep data. The blue and orange colors represent 2 clusters separated by k-means for each quantile. The respective yellow histograms show the distributed cosine distance for each individual pre-transient. Note that for both datasets a dissociation in the RBM weights can be better observed in the bigger transient groups. **(B)** Spatial distribution of activity corresponding to the 2 k-means clusters in the sixth quartile RBM weights. **(C)** Spatiotemporal evolution of the transients triggered by RS or SS modes. The panel shows the average temporal propagation profile (blue: transient start, red: transient end) for each quantile.

In both urethane anesthesia and natural sleep groups, activity preceding large transients is found to be concentrated either in the medial-posterior portion of the cortex, around the RS, or laterally around the SS. Thus, we can conclude that large transients are triggered by small and focused activity bouts and also that such pre-activations are mostly confined to two well-defined cortical regions (Figure 4 B). While smaller events happening to originate in the RS or SS area remained largely confined to the same area for their entire duration, larger events have a growing chance of ‘spilling over’ (Figure 4 C, S4D). Both RS-led and SS-led transients of large size are found to involve different cortical networks over their evolution, generally terminating in the central area of the imaged cortical surface, or even invading the opposite network (so RS activation eventually leading to an activation of the SS network, and vice versa) (Video 2 and Video 3). These findings highlight the presence of two, largely independent cortical networks, one centered around the RS cortex, the other around the SS region (Zingg et al., 2014).

### Neocortical transients preferentially correlate with slow gamma activity in the hippocampus

We then considered what the functional repercussions of transient activity and its structure can be. In particular, cortical sleep activity has been connected to interactions with the hippocampus, possibly supporting memory consolidation (Frankland et al., 2004; McClelland et al., 1995).

In order to explore this question, we analyzed the relationship between global cortical transients and hippocampal local field potentials (LFP). We concentrated on two frequency bands that have been associated to cortico-hippocampal communication, Gamma (20-80 Hz), involved in entorhinal-hippocampal and prefrontal-hippocampal interactions (Bragin et al., 1995; Colgin et al., 2009; Ferraris et al., 2018), and Ripple (150-250 Hz), a component of the Sharp-Wave Ripple complex (SWR) thought to be a main carrier of hippocampal input to the cortex (Battaglia et al., 2004; Sirota et al., 2003).

By averaging the hippocampal CA1 LFP in coincidence with neocortical transients, separately for events belonging to each of the 6 size quantiles defined above, we observed different patterns in the distribution of LFP power density with increasing transient size (Figure 5A). The frequency bands of interests, Gamma and Ripple, appeared to be prominent in the hippocampal LFP spectrum and modulated by the transient size. Indeed, power in both frequency bands showed a significant correlation with simultaneous transient size (Spearman Correlation, Urethane Gamma and Ripple R=1, p=0.001, Sleep Gamma R=1, p=0.001, Ripple R=0.7 p=0.05, Figure 5B). The two bands appeared nevertheless to follow different patterns, with Gamma power presenting a linear increase with transient size, while ripple power following a sigmoid-like trend (indeed ripple power correlation in sleep was only marginally significant, p=0.05).

**Figure 5:**
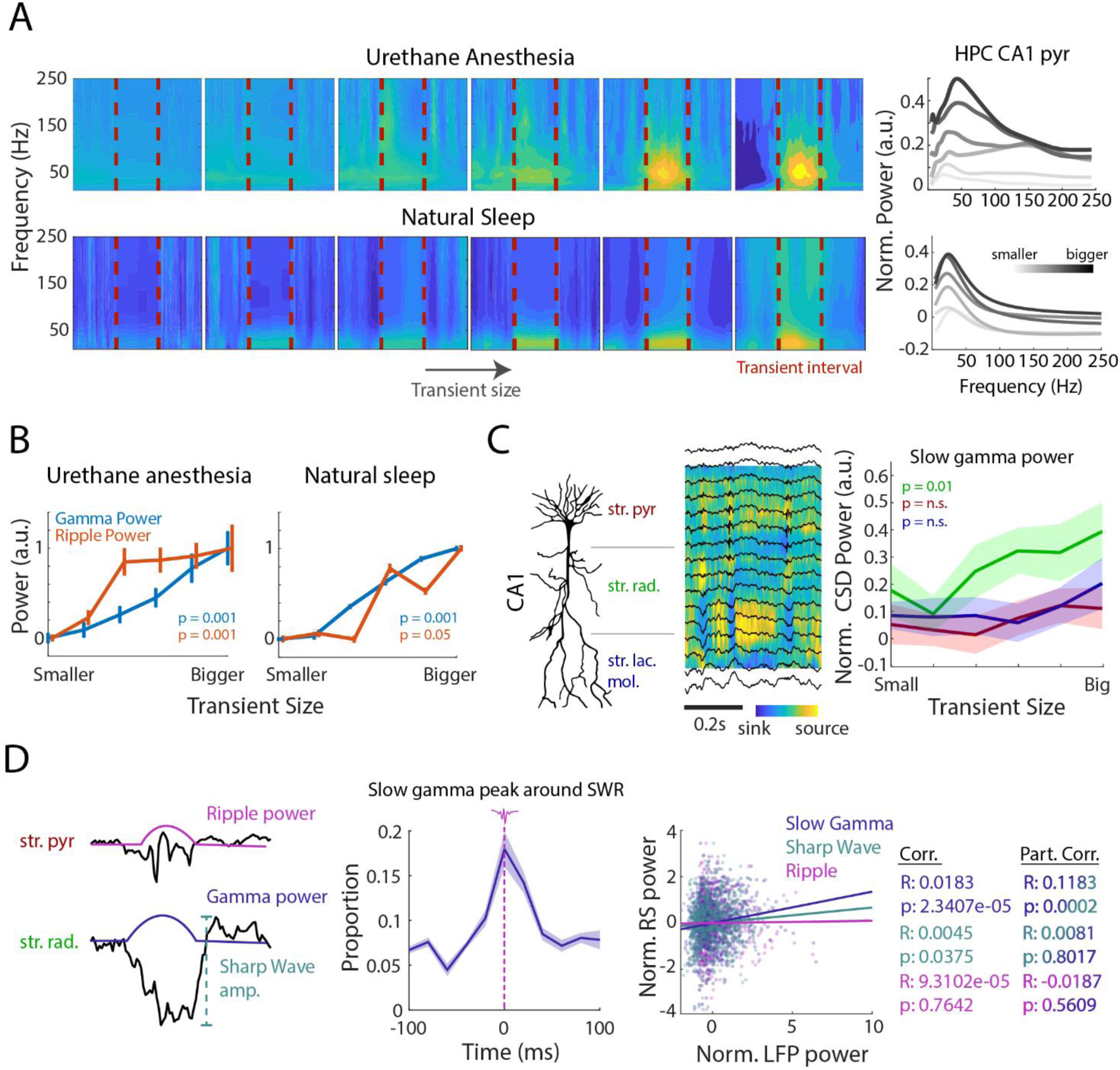
Neocortical transient’s interactions with hippocampal Slow Gamma and SWR. **(A)** CA1 LFP spectrogram (left) and power spectra (right) during transients in the six quantiles in urethane anesthesia and natural sleep data, averaged across animals. Note that the presence of Gamma activity during the transients is proportional to the transient size (a.u. = arbitrary unit). **(B)** Averaged Gamma (20-80 Hz) and Ripple (150-250 Hz) power normalized between 0 and 1 across animals in different transient sizes. (Sperman’s rank correlation coefficient, n = 5 animals). **(C)** (left) depiction of CA1 layers overlayed on a pyramidal neuron silhouette. (center) Layer-resolved LFP (traces) and CSD (color image) from the CA1 layers (right) Normalized CSD power for Slow Gamma (20-40 Hz) in *stratum pyramidale, stratum radiatum* and *stratum lacunosum moleculare* as a function of transient size during natural sleep. A channel in each CA1 layer was selected to calculate the normalized power. The line and respective shadowed area represent the average across animal and SEM (Significance: Spearman one-tailed correlation). **(D)** Interaction between hippocampal LFP and RS activation during SWR. (left) Diagram showing the definition of the three measures (Ripple power, Sharp Wave amplitude and Slow Gamma power) for which we compute correlations with RS activity. (center) Event-triggered average of Slow Gamma peaks centered on ripples. (right) Mean normalized RS activity around the SWR (−100 to 100 ms) (y-axis) against Slow Gamma, Ripple power and the Sharp Wave normalized amplitude (x-axis) in *radiatum* during the Ripples. (right) Correlation of Slow Gamma-RS partialized for Sharp Wave amplitude, Sharp Wave-RS partialized for Slow Gamma amplitude and Ripple-RS partialized for Slow Gamma amplitude.

To further characterize interactions between hippocampal Gamma and cortical activity, in the natural sleep group we looked at layer-resolved signals from linear silicon probe recordings to distinguish between Gamma sources at different depths in CA1. Based on existing literature (Fernández-Ruiz et al., 2017; Lopes-Dos-Santos et al., 2018; Schomburg et al., 2014), distinct Gamma power were computed from the subsequent source area: Slow Gamma (20-50 Hz, *stratum radiatum*), Medium Gamma (60-90 Hz, *stratum lacnosum moleculare*) and High gamma (100-140 Hz, *stratum pyramidale*). We observed that mostly Slow Gamma showed a modulation with the transient size (Figure S5 A). In order to isolate any effect of the volume conduction, we computed Gamma power of the Current Source Density (CSD) signal across different transient sizes. Using CSD signal, we compared Slow Gamma activity obtained from different hippocampal layers. The result confirmed Slow Gamma modulation to be stronger in the *stratum radiatum* (Figure 5 C; Spearman Correlation, R=0.8 p=0.01).

Thus, hippocampal Gamma appears to associated to global cortical activation to at least similar a similar extent as SWR, that are considered the hallmark of cortico-hippocampal interactions.

### Slow gamma accounts for correlations between SWR and cortical activations

As previously reported, SWR events are associated to slow oscillations in the cortex (Battaglia et al., 2004; Siapas and Wilson, 1998), and more recently they have been found to be especially correlated to a network centered around Retrosplenial Cortex (Kaplan et al., 2016; Karimi Abadchi et al., 2020; Nitzan et al., 2020; Opalka et al., 2020). To establish the relevance of SWR events for cortical excitation, we replicated in both urethane anesthesia and natural sleep datasets the approach used by Javad Karimi (Karimi Abadchi et al., 2020). As shown in those studies, we also find that a co-occurrence measure of correlation between SWR events and cortical activation identifies RS as the main hot-spot (Figure S5 B-C). But which feature of SWRs complex is mainly responsible for such relationship? Indeed, SWRs and Slow Gamma events appear to be highly correlated in time (Carr et al., 2012; Oliva et al., 2018) (Figure 5 D), causing their contribution to be potentially intermingled. To answer this question, we considered three SWR properties: Ripple power component, Slow Gamma power component and Sharp Wave amplitude. Indeed, we find that while Ripple power does not correlate with RS activation, the magnitude of the Sharp Wave component of SWR events does show a significant correlation with the degree of RS activation (Correlations: Ripple/RS - R: 9.3102e-5 and p: 0.7642. Sharp wave/RS – R: 0.0045 and p: 0.0375). It is nevertheless the Slow Gamma contribution to the SWR event that appears to have the strongest relationship with RS activity (Correlations: Slow gamma/RS – R: 0.0183 and p: 2.3407e-5). Remarkably, Sharp Wave-to-RS correlation is not significant anymore after controlling for the simultaneous Slow gamma power by partial correlation analysis (Partialized for Slow gamma: Ripple/RS - R: −0.0187 and p: 0.5609. Sharp wave/RS – R: 0.0081 and p: 0.8017). Conversely, the Slow gamma-to-RS correlation remains significant after controlling for the simultaneous Sharp Wave magnitude (partialized for Sharp Wave amplitude: Slow Gamma/RS - R: −0.1183 and p: 0.0002). This set of analysis suggests that at least a portion of the reported dependence of cortical activation on SWR occurrence might be ascribed to the Gamma component present in the latter.

### Transients triggered by RS activation shape Gamma, but not SWR in the Hippocampus

We then asked how our classification of neocortical transients fits into this emerging picture of cortico-hippocampal interactions. We thus repeated the previous analysis using RS-led and SS-led transients separately. Visualizing the average LFP power for different sets of events already makes evident a stark differentiation between the transient families (Figure 6A). RS-led events appear to elicit a much stronger response in the hippocampus, both under urethane anesthesia and in natural sleep groups. By quantifying this effect, we find it to be specific to the Gamma band of the power spectrum (Figure 6 B, Figure S6 B; two-way ANOVA. Urethane anesthesia: Gamma+RS/SS: bins 1-4, p<0.05. Ripple+RS/SS: n.s. Natural sleep: Gamma+RS/SS: bins 4, p<0.05. Ripple+RS/SS: n.s.). Only SS-led transients of major size are found to elicit a comparable Gamma activation in the hippocampus. We speculate that this might possibly happen as a consequence of the transient-spilling previously described, in which case an initially SS-triggered event would eventually result in a delayed RS activation. On the other hand, Ripple power turned out to be largely independent of the transient identity, consistently with our previous analysis on the SWR-RS relationship.

**Figure 6:**
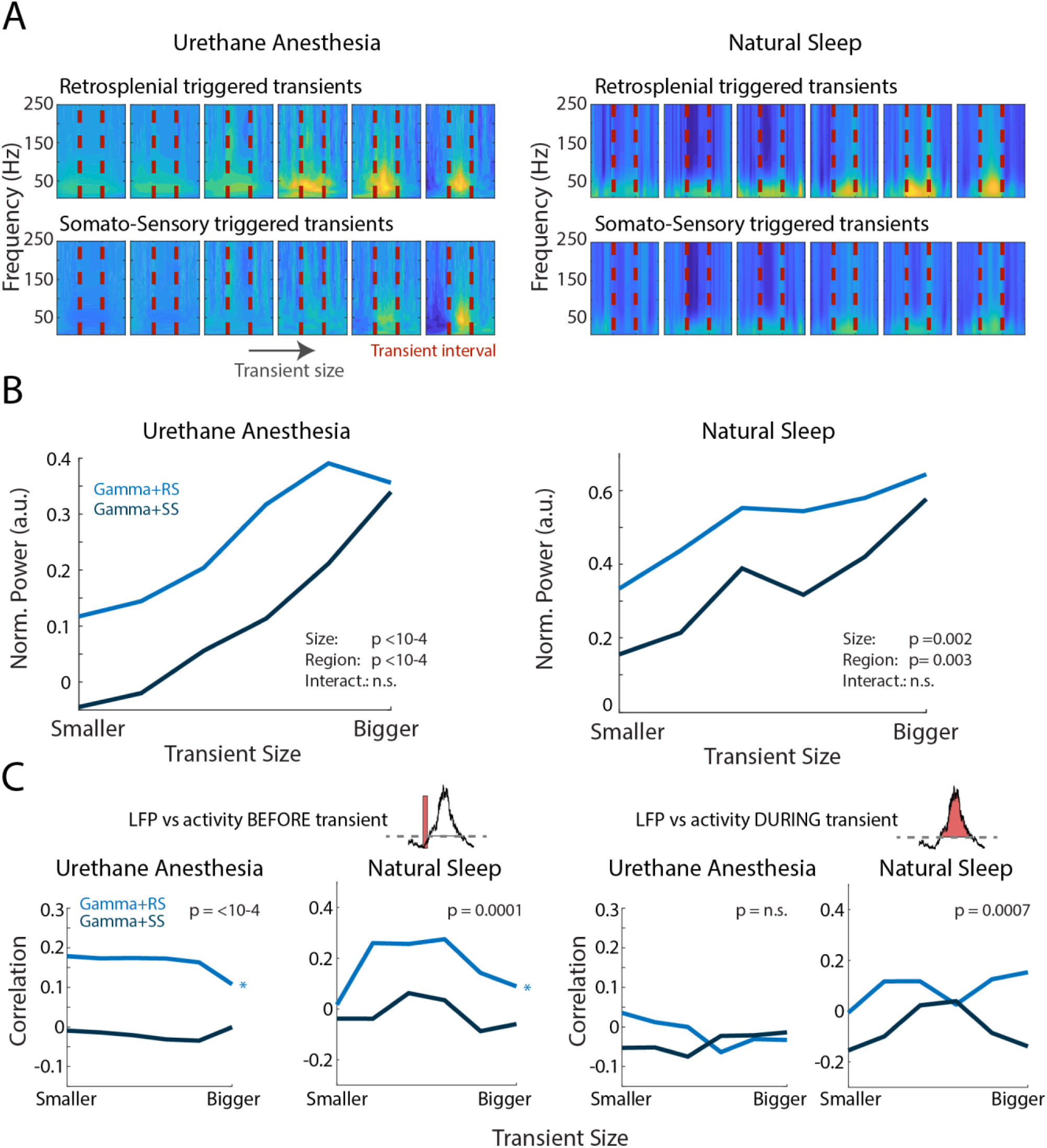
Neocortical transients triggered by RS influence gamma power in the hippocampus. **(A)** Interpolated CA1 spectrogram during the RS and SS triggered transient intervals across different sizes in urethane anesthesia and natural sleep data. **(B)** Normalized power of Gamma (20-80 Hz) for transients triggered by RS and SS across size (*p<0.05; two-way ANOVA). **(C)** Correlation of Gamma with RS or SS triggered events within the same block sizes. The left panel is the correlation between the hippocampal CA1 LFP and the activity of the period before RS and SS transients. The left right is the correlation between the hippocampal CA1 LFP and the activity of the interval during RS and SS transients.

This interaction can be further specified by comparing the magnitude of the RS cortex pre-activation with the elicited power in the Gamma or Ripple band (Figure 6 C, Figure S6 B). By computing the correlation within groups of events of comparable size we find that RS pre-activation scales consistently with the degree of Gamma engagement, and not the Ripple one, during the following cortical transient (p <0.05; two-way ANOVA). This role of RS cortex is time-specific: using its average activity during the transient, thus during a period of cortical excitation, results in no correlation with hippocampal Gamma.

### Precise, bi-directional temporal relationships between cortical and hippocampal activations are orchestrated by RS

Finally, we investigated the temporal structure of the cortico-hippocampal interactions outlined above. We first computed the cross-correlation between the temporal evolution of hippocampal power in either the Ripple or Gamma band and the degree of activation of different portions of the cortical surface (Figure 7 A). Importantly, we used partial correlations, that is, we computed the correlation between pixel activation and Gamma power controlling for Ripple power, and vice versa (cortical activity vs. Ripple controlling for Gamma). Confirming our previous findings, correlations between Gamma and cortex are much stronger than those involving Ripples. Both in urethane anesthesia and natural sleep recordings correlations appear to follow a precise spatiotemporal pattern. Positive correlations with Gamma power tend to appear at negative time shifts, thus preceding the peak of the Gamma power. Also, they are initially localized in areas close to the midline and they mostly involve cortical areas distributed either medially or posteriorly, overlapping with the default mode network. In contrast Ripple power is associated with a much weaker coherence in cortical activation and with a higher degree of symmetry around its peak.

**Figure 7:**
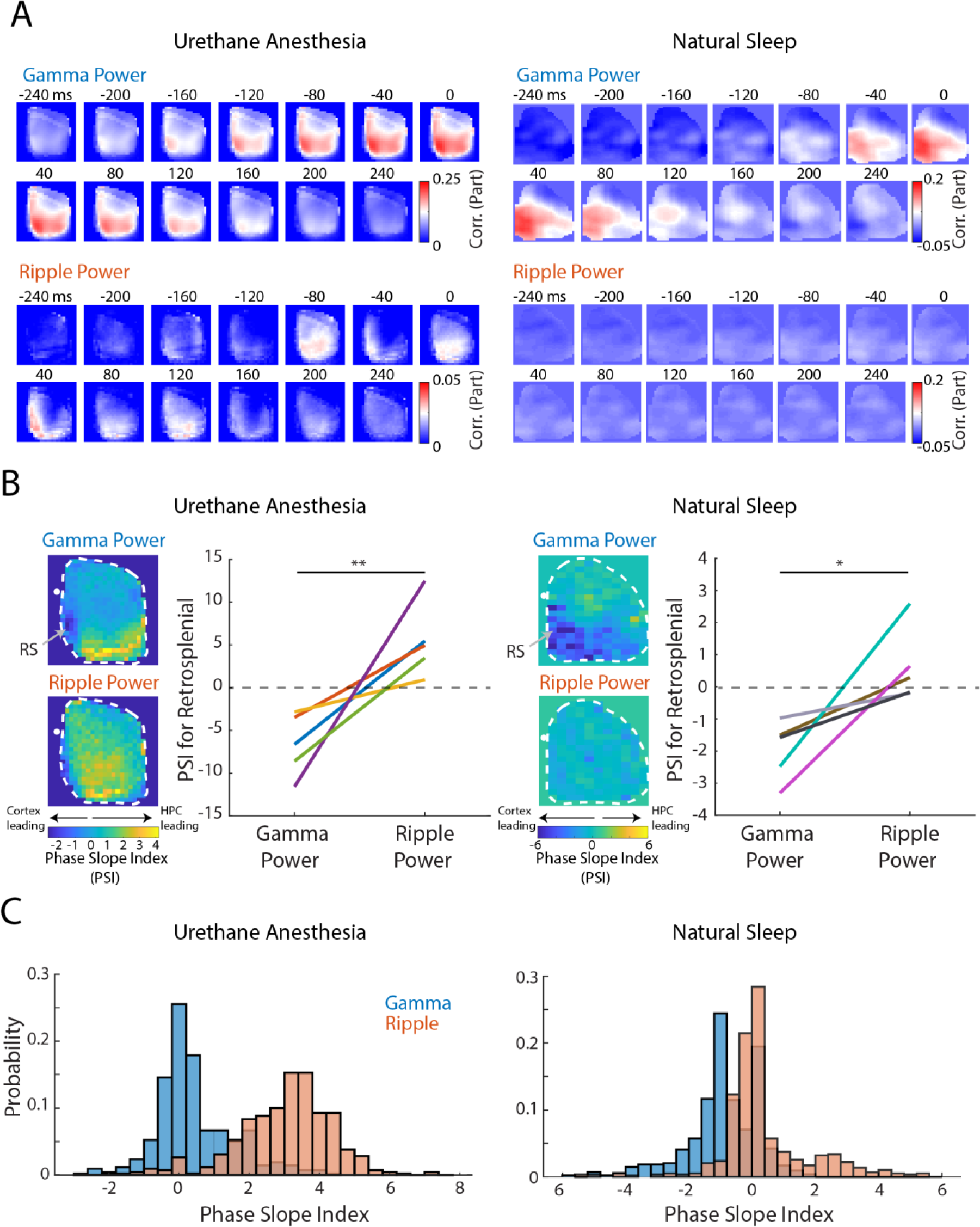
Bidirectional interactions between hippocampus and neocortex. **(A)** Partial correlation between hippocampal Gamma power versus the optical temporal signal for each pixel partialized for Ripple power and partial correlation between hippocampal Ripple power versus the optical temporal signal in each pixel partialized for Gamma power. Correlations were calculated from z-scored data. **(B)** Phase slope index for normalized Gamma or Ripple power versus the optical temporal signal in each pixel. The white dot represents bregma. The gray arrows indicate the position of Retrosplenial cortex, which presents a temporal modulation, in Gamma, but not in Ripple, from the cortex to the hippocampus (*p<0.05, and **p<0.005; one-sample t-test against zero; n=5 animals for the urethane and n=5 for the natural sleep group represented in different colors). **(C)** Phase Slope Index scores for all the neocortical imaged pixels calculated for Gamma and Ripple powers.

Not only then hippocampal engagement is preferentially associated to a network organized around the RS cortex, but such engagement appears to be initiated by the cortex. This view is confirmed when computing the Phase Slope Index (Nolte et al., 2010), a measure of directed causality, between cortical activation at different sites and hippocampal LFP instantaneous power. Conversely, Ripple power leads a large swath of the cortex with little spatial specificity. Directionality scores are in fact higher than those obtained from Gamma power (Figure 7 B). Strikingly, the RS region present an extreme version of this tendency (one-sample t-test, between Gamma and Ripple for urethane anesthesia: p<0.005. Gamma and Ripple for natural sleep: p<0.05. Comparing to zero, urethane anesthesia: Gamma p<0.005, Ripple p<0.05 and natural sleep: Gamma p<0.005, Ripple n.s.). Interaction directionality scores computed from this area in fact present an almost complete switch between Gamma and Ripple power band: they are consistently negative in the former case and distributed around zero in the latter. Thus, RS cortex alone (in the range of cortical areas we were able to image), is responsible for leading the hippocampal Gamma events, while Ripple events are associated with a more spatially generalized flow of information from the hippocampus to the cortex (Figure 7 C).

## Discussion

### Quasi-critical nature of cortical spontaneous activity during sleep

The transient, fluctuating nature of cortical activity during NREM sleep has been known for a long time (Steriade et al., 1993b). Slow oscillations that can be detected from cortical activity with EEG and other measuring modalities are the global manifestation of UP/DOWN fluctuations in membrane potentials widespread in cortical neurons. “UP” states are the result of recurrent excitatory feedback in cortical modules (Shu et al., 2006; Steriade, 2001; Waters and Helmchen, 2006), and propagate in wave-like fashion, potentially to very far location in the cortex (Amzica and Steriade, 1995b; Huber et al., 2004; Mohajerani et al., 2010). UP states are abruptly terminated due to a variety of causes (Holcman and Tsodyks, 2006; Luthi and McCormick, 1998; Massimini and Amzica, 2001), and that termination may happen after excitation has spread to arbitrarily large cortical territories (up to the entire cortex). In fact, computational models (Levina et al., 2007; Millman et al., 2010) showed that recurrent excitation and short-term synaptic depression (a candidate cause for UP state termination) are sufficient ingredients to obtain a (quasi)-critical distribution of UP state sizes. Similar to what described during recovery from anesthesia (Scott et al., 2014), this is what we show here, in NREM-like urethane anesthesia and in natural NREM sleep, with Voltage imaging and ECoG.

While anesthesia Voltage imaging data show an excess of very large events (essentially covering the entire imaged portion of the cortex), in natural sleep there is a tight fit with a power-law, signature of critical processes (Figure 2A).

We suggest that critical dynamics could play a very important role in memory processes, which should be seen in the light of systems consolidation theory. Any complex long-term memory will involve information stored at multiple, far away sites. The hippocampus is thought to generate rapidly constructed and readily retrievable index representations, which may ignite these whole-cortex representations. Yet, this poses the question of how the limited hippocampal input, propagated through relatively sparse long-range pathways may accomplish this feat. Critical dynamics is a regime of enhanced long-range correlations, yielding increased information storage (Boedecker et al., 2012; Wilting and Priesemann, 2019), and transmission (Shriki and Yellin, 2016). As such, criticality facilitates the cortex-wide spread of hippocampal signals, and provides a basis for coherent cortico-hippocampal representations, beyond the few hub areas with a close link to the hippocampus (such as prefrontal or entorhinal cortex). Indeed, coherent replay has been observed between the hippocampus and sensory areas with only an indirect, polysynaptic link with it, such as auditory (Rothschild et al., 2017) and visual cortex (Ji and Wilson, 2007).

Furthermore, cortical transients may be self-ignited, and correspond to spontaneous processing and – during sleep – memory retrieval. Initial evidence that this may happen comes from magnetoencephalography studies with human subjects (Higgins et al., 2020; Liu et al., 2019) showing spontaneous replay events in the cortex. A theoretical perspective (McClelland et al., 1995, 2020) is that cortically-generated replays retrieve older memories, and would be interspersed with hippocampus-triggered replay of recent memories, an arrangement that would protect memory from catastrophic interference, the sudden loss of all memories due to exceedingly fast encoding of new memories.

### Cortical transients activate anatomically-determined networks

A purely statistical description in terms of transient sizes misses however the highly structured character of cortical connectivity and activity. Here, we show that multi-scale transients are organized around networks. Our data-driven analysis demonstrates (Figure 3), in a highly consistent fashion across two preparations, that small transient activations mostly remain confined to one of three networks, centered respectively on medial areas, anterior sensorimotor cortices, and lateral cortex.

Enticingly, there is an excellent correspondence of the activity arrangement stemming out of our analysis with the organization delineated with a network-theoretic analysis based on a massive amount of anatomical projection experiments. In the paper by Zingg et al. (2014), a clustering of cortical connections is described into four networks. Our ‘RBM 2’ (Figure 3) component strongly overlaps with the ‘sensorimotor network’ in that study, combining somatosensory and primary motor cortices. A set of ‘medial subnetworks’ highlighted by Zingg et al. overlaps with our ‘RBM 3’ component. That network, both in our data and in the anatomical characterization is centered around the retrosplenial cortex, and overlaps with the Default Mode Network as defined in the mouse brain (Stafford et al., 2014). Our ‘RBM 1’ component contains part of what Zingg et al. call the ‘posterior temporal lateral’ network, which includes structures such as perirhinal and TeA cortices, that are major players in the up and down streams of highly processed sensory information into the medial temporal lobe and eventually the hippocampus. An obvious limitation of our study is that, by imaging only dorsal cortices, we miss medial bank and ventral cortices, and therefore much of the lateral network in particular, as well as DMN areas, such as medial prefrontal cortex. Still, our data provide evidence that anatomical connectivity shapes the transient constituents of spontaneous activity, in a fashion that likely affects its function.

Larger transients eventually invade a large portion of the recorded cortex, and are necessarily less clustered. Yet, if we look at the cortical activity immediately preceding them, we see that the activity that ignites them is still circumscribed to one of the networks described above. Our ‘RBM 1’ component (Figure 4) shows activation confined to retrosplenial cortex and other areas in the DMN, whereas ‘RBM 2’ shows transients with a sensorimotor origin. It is possible, due to the limitations in our field of view, that we missed transients initiated, for example, in the lateral networks, but our results highlight how states of generalized cortical activation (potentially carrying complex memories) may be started in different networks, which would contribute memories of a different nature (e.g. procedural vs. declarative) or age.

### Bi-directional interactions between cortical networks and hippocampal activity

The ‘standard model’ of systems consolidation emphasizes how hippocampal activity, with its rich content in replayed information, may affect spontaneous activity in the neocortex, coordinating the memory consolidation process (Frankland and Bontempi, 2005; McClelland et al., 1995). The hippocampal sharp wave-ripple (SWR) complex is considered to be the main conduit for this information transfer, for two reasons. First, because they coincide with the highest density of replay (Kudrimoti et al., 1999) and second because they correlate with a widespread increase in cortical firing (Battaglia et al., 2004; Logothetis et al., 2012; Mohajerani et al., 2010; Sirota et al., 2003) and replay (Ji and Wilson, 2007; Peyrache et al., 2009). The Default Mode Network (DMN) is found to be a preferential recipient of that input (Kaplan et al., 2016), probably at least partly mediated by direct afferents for example to retrosplenial cortex (Nitzan et al., 2020). The majority of contemporary computational models of memory consolidation assume this one-way flow of information during sleep (Kali and Dayan, 2004; Singh et al., 2022), from the hippocampal source to the neocortical target.

Our imaging data (Figure 5) are consistent with this view: hippocampal ripple oscillation power correlates with small and large transients (Figure 5B) with similar intensity. PSI, a pseudo-causality analysis, suggests that ripple power predicts cortical transients (Figure 7). Thus, SWRs and potentially replay, are able to elicit widespread neocortical UP states at multiple spatial scales. Yet, a surprisingly strong link was observed between hippocampal Slow Gamma and cortical transients. Whenever a comparison was possible, for example by partial correlation analysis, we found that the Slow Gamma link dominated the association between Ripples and neocortical activity (Figure 6B and S6). PSI for Slow Gamma shows an opposite pseudo-causality flow, from neocortical activity to hippocampal Slow Gamma. It is known that hippocampal activity is affected by cortical UP-DOWN state fluctuations (Isomura et al., 2006; Sirota and Buzsaki, 2005), even at the level of resting membrane potential (Hahn et al., 2006). Our results identify the global cortical activity patterns that may affect the hippocampus and highlight one activity mode, Slow Gamma oscillations, as a candidate mediator of cortical influences.

During wakefulness, slow gamma oscillations fluctuate greatly (Bieri et al., 2014; Cabral et al., 2014; Lasztóczi and Klausberger, 2016). The instantaneous balance between Slow Gamma and medium gamma may reflect the momentary prevalence of respectively CA3 vs. entorhinal inputs into CA1 (Bragin et al., 1995; Colgin et al., 2009). Correspondingly, a shift between memory retrieval, predictive hippocampal activity and registration and storage of novel cortical inputs may take place (Guardamagna et al., 2021). During sleep, SWR events associated to higher levels of Slow Gamma contain increased replay (Carr et al., 2012). Together, these results point at slow gamma as an index of self-initiated hippocampal processing, reflecting previous experience.

Our data may be explained by positing that cortical transients, by depolarizing neurons, may bias the hippocampal networks into a slow-gamma rich state. It is also likely in our view that the information content of the neocortical input will modulate which activity patterns will be retrieved (or generated) by the hippocampus. This interpretation is consistent with a bi-directional interaction between cortex and hippocampus (Figure 8) during sleep. A possibility is that the cortex will bias the hippocampus into providing the most needed content at any point in time to carry on the memory consolidation process (Battaglia et al., 2011; Sirota and Buzsaki, 2005). A more radical stance, which would have to be explored experimentally and computationally is that neocortex and hippocampus act in the sleep state as a single network, with transient activations that may be initiated in several points in the network and propagate at multiple scales.

**Figure 8:**
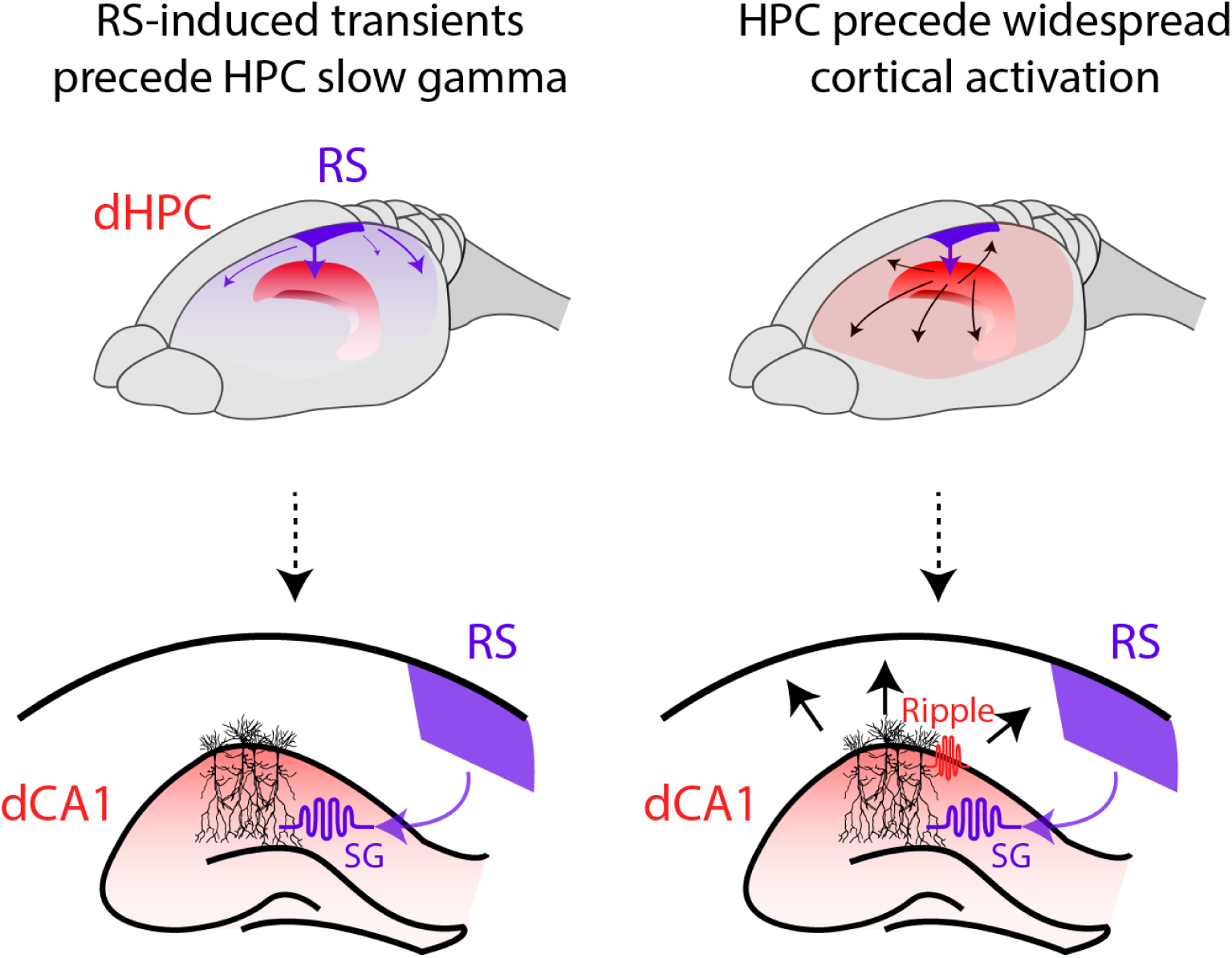
Schematic of the hypothesized bi-directional interaction between the RS network and the hippocampus.

### Retrosplenial cortex and the DMN preferentially engage hippocampal activity

Importantly, not all neocortical transients interact with the hippocampus in equal manner (Karimi Abadchi et al., 2020). The key distinction apparent from our analysis is between transients initiated in the medial network, with a focus on retrosplenial cortex, and those initiated in the sensorimotor network (Figure 4). The former is related to a significantly larger increase in slow gamma power in the hippocampus (Figure 5A-B) and with a tighter correlation (Figure 5C) than the latter. Within the medial network, the role of retrosplenial cortex stands out: not only is RS the hotspot for initiating large transients (“RBM 1” in Figure 4B), but it is also the area whose activity most markedly predicts increases in hippocampal slow gamma. The rest of the medial network (and of the recorded neocortex) ignites at a later moment (Figure 7A) following hippocampal activation (Figure 7B). Thus, the medial network activation may activate independently of the hippocampus. In humans, the DMN, largely overlapping with the medial network observed here, has been found to activate in correspondence with replay bursts (Higgins et al., 2021), suggesting that it plays a distinct role in memory retrieval and recollection. Because the retrosplenial cortex, also a hub of the DMN, has been found to be a hub for remote memory retrieval (Cowansage et al., 2014), it is possible that the large transients initiating there do represent replay of older memories, to be interleaved with new memory replay, initiated by the hippocampus, a set up that is thought to be beneficial for memory consolidation. We speculate also that the RS-initiated replay may provide the semantic context for reprocessing in the hippocampus.

In sum, we presented a novel view on the global architecture of cortico-hippocampal interactions, which opens new theoretical perspectives on the mechanisms of memory consolidation. Further investigation of these mechanisms will require, among other things, understanding how hippocampal ensembles are affected by cortical influences, in a view emphasizing two-way exchanges between these two brain areas key for memory and cognition.

## Materials and Methods

### Experimental model and subject details

The experiments for the urethane group were carried out in five C57/Bl6 adult mice (age 2 to 4 month), male and between 20 and 30 g for acute VSD imaging. In the end of the experiments, they received a pulse of electrical or photoelectric stimulation in the hindlimb, forelimb and eye for later mapping of the cortical topography. The animals were housed in standard plastic boxes shared with other animals under the conditions of 12-h light/dark cycle. The animals had ad libitum access to water and food all the time. All the precedents indicated by the Canadian Council on Animal Care (CCAC) were followed and the protocols of animal experimentation were approved by the Animal Welfare Committee of the University of Lethbridge.

The natural sleep experiments were performed in five CaMK2A-tTA;tetO-chiVSFP mice (male and age 3 to 6 month), between 25 and 35 g for acute VSI imaging. The animals were group housed until the surgery day, and then individually housed after to prevent damage to the implants. They had continuous ad libitum access to water and food and were on a 12-h light/dark cycle. All the experiments in sleep conditions were performed during the light period. This experimental procedure was approved by the Central Commissie Dierproeven (CCD) and by the Radboud University Animal Welfare Board and conducted in accordance with the Experiments on Animals Act and the European Directive 2010/63/EU on animal research.

### Surgery

#### Surgical preparation for urethane recording

Mice were initially anesthetized with 3% isoflurane at 100% O_2_ concentration. Animals were then fixed in a stereotaxic apparatus and maintained under isoflurane anesthesia at 0.5-1% by a nasal mask. The body temperature was maintained at 37 °C with a heating pad regulated by a feedback thermistor. A dose of 80 μg of dexamethasone (intramuscularly) was injected to relieve pain, followed by 50 μl of lidocaine at 2% topically in the area where the incision was to be made. For hippocampal registration, 2 Teflon-coated stainless-steel wires (277 μm diameter) were implanted on the pial surface of the cerebellum as ground and in the hippocampus (−4.3 mm AP and +2.4 mm ML, +1.8 mm DV at a 60-degree angle). The electrodes were carefully fixed to the skull with super glue and acrylic resin.

Subsequently, a tracheostomy was performed to facilitate the respiration of the animal. A half hemisphere craniotomy window was made for skull removal. After the craniotomy, the cortical surface was carefully covered with 4- (2-hydroxyethyl) -1-piperazineethanesulfonic acid (HEPES) - buffered saline solution (1 mg ml^− 1^) and applied to the exposed cortex to maintain hydration. At the end of all these procedures, the animals were anesthetized with 15% urethane (1250 mg / kg) and their isoflurane masks were removed.

#### Surgical preparation for natural sleep recording

Animals were anesthetized with 2% isoflurane at 100% O_2_ concentration and then fixed in a stereotaxic apparatus and maintained under isoflurane at 0.5-1.5% by a nasal mask. The body temperature was maintained at 37 °C with a heating pad regulated by a feedback thermistor. A subcutaneous dose of 5 mg/kg of carprofen was injected at the onset of the anesthesia, followed by 50 μl of lidocaine at 2% (subcutaneous) through the scalp. Isoflurane and Oxygen level were observed and controlled during the surgery to maintain the breathing rate at 0.5-1.5 Hz. The skull was then exposed and a skull screw implanted above the left cerebellum to provide common reference and ground to the silicon probe. Afterwards, in order to better fix the head-plate implant, 2 extra skull screws were also implanted on the left hemisphere. The exposed skull in the right hemisphere was then thinned in order to remove small skull capillaries to improve the later voltage imaging recording. For CA1 laminar recordings, we performed a craniotomy on the pial surface of the right cerebellum (−4.3 mm AP and +2.4 mm ML) and a 16 channels silicon probe with an inter-site pitch of 50 μm (E16_R_50_S1_L10, Atlas Neuroengineering, Belgium) was inserted in the intermediate hippocampus (+2.1 mm DV at a 57-degree angle towards the back). The electrodes were carefully fixed to the skull with cyanoacrylate and acrylic resin. To finish we placed a 3D-printed custom-designed head-plate with a wide field view of the right hemisphere fixing it on the skull with acrylic cement (Super-Bond C&B). After the surgery all animals were given at least 3 days before beginning the habituation in the setup. See (Pedrosa and Battaglia, 2021) for a detailed description of the protocol.

### Voltage sensitive dye imaging (urethane group)

After the surgery, the voltage-sensitive preparation dye solution was set up based on previous work from our group(Bermudez-Contreras et al., 2018; Karimi Abadchi et al., 2020). Initially the voltage dye RH1691 (Optical Imaging, New York, NY), was dissolved in HEPES-buffered saline solution (0.5 mg/ml) and applied for 30-50 min in the exposed brain. To minimize the movement artifacts due to the respiration and heartbeat, the stained brain was covered with 1.5% agarose made in HEPES-buffered saline and sealed with a glass coverslip. For image recording we used a charge-coupled device (CCD; 1M60 Pantera, Dalsa, Waterloo, ON) and an EPIX E8 frame grabber with XCAP 3.7 imaging software (EPIX, Inc, Buffalo Grove, IL) to record 12-bit images every 5 ms (200 Hz). To excite the dye, we used a red LED (Luxeon K2, 627 nm center) together with excitation filters of 630 ± 15 nm. Reflected VSD fluorescence was filtered using a 673 to 703 nm bandpass optical filter (Semrock, New York, NY). The optical images were acquired through a macroscope composed of front-to-front lenses (8.6 × 8.6 mm field of view, 67 µm per pixel). To reduce potential artifacts caused by the presence of neocortical blood vessels, we focused the lens into a depth of ∼1 mm from the cortex surface. All the experiments were recorded in a duration ranging from one to two hours. In order to identify the topographic map of the neocortical areas, the voltage optical signal was also recorded from the neocortex in response to different sensory stimuli in the periphery, as described previously (Mohajerani et al., 2013). Thereby, the coordination for the primary sensory areas (HLS1, FLS1, V1 and A1) and secondary somatosensory areas (HLS2 and FLS2) were identified. Based on the evoked activity of these areas and bregma coordination, the relative locations of additional associational areas were estimated using stereotaxic coordinates (RS, ptA, M2/AC) together with others areas as: V2L (lateral secondary visual cortex), M1 (primary motor cortex), BSC (Barrel cortex) and S1/S2 (primary and secondary sensory areas).

The hippocampal LFP was recorded from teflon coated 50.8 µm stainless steel wires (A-M Systems). For that, based on previous work (Karimi Abadchi et al., 2020; Pedrosa and Battaglia, 2021), a craniotomy was performed on the right hemisphere skull about +2.4 mm and tangent to the posterior side of the occipital suture. Next, the electrode was placed at a 57-degree angle in relation to the vertical axis and was gradually lowered through the brain. During lowering, the signal was monitored visually and audibly until a large increase in the MUA was observed close to the expected coordinate for the pyramidal layer in dorsal CA1 (depth = ∼1.75 mm). After setting, we glued the electrode on the skull using Krazy Glue and dental cement.

### Genetically encoded voltage indicator imaging (natural sleep group)

The GEVI used, chiVSFP, reports membrane voltage by decease and increase in yellow (mCitrine) and red (mKate2) fluorescence. To image these signals, a epifluorescence macroscope setup equipped with Leica PlanAPO1.6 lens based on previous works was used (Scimedia, Costa Mesa, CA) (Akemann et al., 2010; Carandini et al., 2015; Ratzlaff and Grinvald, 1991). Excitation light was provided by a 150 W halogen lamp (Moritex, Brain Vision). The optical filters (Semrock) used were: mCitrine excitation 500/24, mCitrine emission FF01-542/27, mKate2 emission BLP01-594R-25 and excitation and detection beam splitters (515LP,580LP). Fluorescence images were aquired with two synchronised sCMOS cameras (PCO edge 4.2) controlled by a TTL external trigger. The images were acquired at 50 Hz in several blocks of 15 minutes at 375 ×213 pixel (spatial resolution of ∼36 um/pixel) and 12-bit digitalization. One of the animals had the images acquired at 25 Hz in blocks of 20 min.

### Hippocampal LFP and Wide-field ECoG recording for urethane anesthesia

The hippocampal LFP recordings were amplified (x 1,000) and filtered (0.1-10000 Hz) using a pre-amplifier Grass A.C. Model P511 (Artisan Technology Group, IL) and digitized in a Digidata 1440 (Molecular Device Inc, CA) data acquisition system at 20 kHz sampling rate.

For the electrophysiological profile of the neocortex, we used a transparent electrocorticography grid. The recording sites on the grid is organized in a matrix 6×5 (spaced in 1 mm between sites) covering a total area of ∼6 mm x 5 mm. To record the neocortical activity, we first grounded the grid to the nose of the animal with an electrode implanted subcutaneously. Then, we carefully set the grid over the right neocortex of the mouse, thus covering the same wide-field recorded in the optical imaging. After, we covered the exposed cortical area with HEPES-buffered saline to maintain hydration. The electrophysiological signals were recorded using 32-channels headstages (RHD2132, Intan Technologies) connected to an Open Ephys recording system. The raw signals were filtered between 0.1 and 7500 Hz, pre-amplified (20x) before digitization at 20 kS/s.

### Hippocampal LFP probe recording for natural sleep

The hippocampal LFP signals were acquired using 16-channels headstages (RHD2132, Intan Technologies) connected to an Open Ephys recording system. The raw signals were filtered between 0.1 and 7500 Hz, pre-amplified (20x) before digitization at 20 kS/s.

### VSD data preprocessing (urethane group)

The VSD raw data recorded was preprocessed by first correcting in Matlab the time-dependent reduction of the fluorescence. For this the time series relative to each pixel was filtered using a zero-phase highpass Chebyshev filter above 0.2 Hz. Then, a baseline signal (F₀) was computed based on the average of all the pixels recorded in each frame. The baseline signal calculated was then subtracted from the filtered signal, and the difference was divided by the baseline values at each time point (F-F₀/F₀ × 100, where F is the filtered signal).

### VSI data preprocessing (natural sleep group)

The voltage signal was calculated from the fluorescence signals as described in previous work (Akemann et al., 2012; Song et al., 2018). Gain equalization, as described in Akemann et al. (2012) minimized vascular artifacts and hemodynamic components in the ratio (R) of mKate to mCitrine fluorescence intensity. The voltage signal was calculated as R-R₀/R₀ × 100. The resultant voltage map was then spatially smoothed (2D Gaussian convolution, 20 × 20 pixels) and temporally filtered (Chebyshev high-pass filter, >0.5 Hz).

### Facial camera and pupil detection

On the left side of the animal’s visual field (at 45 degrees from the center and 12 cm of distance), a camera (Basler aca1920-150um) attached to 2 infrared LEDs filmed at 20 Hz the facial action during the sleep periods. Offline, pupil contour was tracked using the DeepLabCut toolbox (Mathis et al., 2018).

### Sleep recording and scoring

After a period of adaptation in which the animal was placed in the head-fixed setup daily for two weeks, natural sleep was recorded during the light cycle. Data were acquired from moment the animal entered in the first REM sleep episode.

Offline, the data was initially classified as putative NREM or REM sleep based initially on the Theta/Delta ratio volume from the CA1 pyramidal layer. The awake periods were then identified based on visual inspection in the behavioral data recorded from the facial camera and an encoder connected to the floor of the animal. REM sleep periods were confirmed based on a reduction in pupil size (Yüzgeç et al., 2018). Lastly, NREM periods were settled as the periods before the onset of the REM sleep until 30s after the animal expresses an awake behavior (walk or move the chin or nose for at least 1 second). In this study, only the states classified as NREM sleep were analyzed.

### SWRs detection

For the SWRs detection we adapted the method used in (Peyrache et al., 2009). Initially, the data was down-sampled to 1 kHz and filtered in the ripple band (150-300 Hz). The root mean square was then smoothed by convolution with a gaussian window of 30 ms. Thereafter, the peaks of the filtered signal in between 3 and 5 s.d. plus the mean were detected. The start and end of the ripple was identified based on the points that crossed 1.5 s.d. before and after the detected points. We took as ripple events the intervals so detected that were longer than 30 ms and shorter than 300 ms. Also, the distance between the end of a ripple and the start of the next had to be at least 100 ms or the events were merged.

### Cortical transient detection

The procedure to identify cortical transients was equivalent for VSD, VSI and ECoG data.

Cortical transients were defined by applying a combination of two thresholds. Each channel signal time-series *s*_*i*_*(t)* was z-scored and then a threshold at 2 std was applied, so that 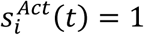 whenever 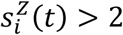 and 0 otherwise. The global state of activation was then computed as the number of channels active at any time 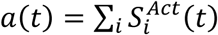. In turn we applied a second threshold to *a(t)*: a transient was taken to be present at time *t* whenever the value of *a(t)* would be above its mean computed over the entire recording session. Individual transient events were then defined as continuous stretches of active time bins between two threshold crossings of a(t), 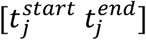. Each transient was then characterized by its 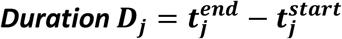 and by its 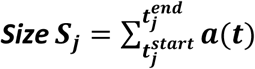. For each recording session transients were grouped into 6 groups based on their size. Size limits were determined by dividing the observed size range into logarithmically spaced intervals.

### Restricted Boltzmann Machine Classification

Cortical transient spatial activation patterns were classified via a Restricted Boltzmann Machine (RBM) procedure (Figure S3 A-E). For each recording session, we first collected all time frames belonging to transients of one size group. This collection of instantaneous activity configurations was used as training inputs for an RBM with n=50 hidden units. Learning was performed in a standard way through momentum gradient descent (Hinton, 2012), over 50 training epochs, with 3 contrastive divergence steps for each input pattern, a learning rate of eta = 0.001 and weight regularization. We trained a separate RBM for each size group and either i) using time frames belonging to identified cortical transients or ii) time frames from the 100 ms preceding the detected onset of each transient in the group. Once completed we collected the weight values resulting from the training procedure. We then looked for the existence of repeating activity patterns in the cortical space by analysing the distribution of the weights over the input bins developed by each hidden unit in the RBM. In particular we were interested in identifying clustering of weight profiles over the hidden units as they would indicate the presence of highly likely activity configurations (Olshausen and Field, 1996). We thus proceeded in two ways, on the one hand we applied multidimensional scaling (non-metric MDS, sammon algorithm) to the set of weight vectors and investigated their distribution after dimensionality reduction. Most-likely number of clusters was assessed by fitting a mixture of Gaussians over this distribution through EM algorithm and varying the number *k* of independent gaussians in the mixture. For each condition and session, the most likely number of clusters was established by comparing results with different k after correction for parameter number using Akaike Information Criterion modified for limited sample size. The presence of actual clustering in the original weights before dimensionality reduction was then confirmed by looking at the distribution of correlations among different weight vectors. A strong bimodal-like shape in the distribution was taken as evidence for the presence of clusters. Significant cluster organization was found for weights obtained from activity of small transients (3 clusters, 4 out of 5 urethane animals and 4 out of 5 natural sleep animals) and for weights obtained from activity preceding large transients (2 clusters, 5 out of 5 urethane animals and 3 out of 5 natural sleep animals). We then used the most-likely cluster number to classify hidden units in *k* classes (k-means clustering). For each class an average weight vector was computed, and used to generate a typical spatial activation pattern for that class. Spatial distributions were highly regular across animals and were therefore averaged to produce activation templates both for the small transient case and for the preceding-large transient one.

### Region-Specific Cortical Transients

To classify transients in either Retrosplenial (RS) or Somatosensory (SS) initiated we computed the overlap between the activity preceding each one of them and the RS and SS activation template (obtained from the RBM classification), respectively. We then took transients with either above-average RS or SS overlap as belonging to one of the two types. Similarly, RBM templates were used to measure the average expression of specific patterns over the duration of transients of different size. The overlap between each template and transient activity was computed for each time bin of a transient, and then averaged across transients after having normalized their duration to the [0 1] range (Figure S4D). Alternatively, a similar time-resolved analysis of the transient evolution was performed using PCA instead of BM-templates. In this case PC analysis was performed over the entire duration of one recording session. The first 10 PCs (>80% variance explained) were then retained and used as a basis for projecting cortical activity during each transient. Average across transients was then performed using the same transient duration normalization step (Figure 4C).

### Phase Slope Index Analysis

We investigated the presence of directionality in the cortico-hippocampal interactions employing the Phase Slope Index analysis introduced by Nolte et al. (Nolte et al., 2010). The analysis is based on measuring the scaling of phase difference at different frequencies between two continuous signals. It returns a score indicating both the direction of the interaction and its strength. We applied this method to test the presence of a ‘causal’ relationship between the raw cortical VS signal and the power of the hippocampal LFP filtered in either the ripple or the slow-gamma band.

## Statistical tests

### Power-Law scaling fits

Scaling exponent was assessed by linear regression of the log transformed data. Quality of the fit was measured by uncertainty on the inferred fit parameters.

### Size effects

To test for transient size correlation a one-tailed Spearman-Rank correlation was used.

### Region differences

2-way ANOVA was run.

## Acknowledgments

We acknowledge support from the European Commission Horizon 2020 program, grants ERC-AdG 833964 “REPLAY_DMN” (to FPB), MSCA ITN 765549 “M-GATE” (to FPB), MSCA Intraeuropean Fellowship 840704 “BrownianReactivation” (to FS and FPB), and from Natural Sciences and Engineering Research Council of Canada 819 (grant no 40352 & 1631465 to MM), Alberta Innovates (to MM), 820 Alberta Prion Research Institute (grant no. 43568 to MM), and Canadian Institute for Health 821 Research (grant no 390930 & 156040 to MM), National Science 822 Foundation (MM), National Institute of Health (1U01NS099573 to TK). The authors also thank J. Sun for surgical assistance, Matteo Guardamagna, Jeroen Bos and Chenchen Song for technical advice, Tjitse van der Molen for help with data analysis, and Christophe Bernard and Loig Kergoat for donating the ECoG grids.

## Contributions

R.P., M.M., F.S. T.K. and F.B. designed the experiments. R.P. and M.N conducted the experiments. R.P., F.S. and F.B. designed the analysis techniques. R.P. and F.S. performed the data analysis. T.K. provided the experimental models R.P., M.M., T.K., F.S. and F.B. wrote and edited the manuscript. F.B. supervised research.

## Supplemental Information

**Figure S1:**
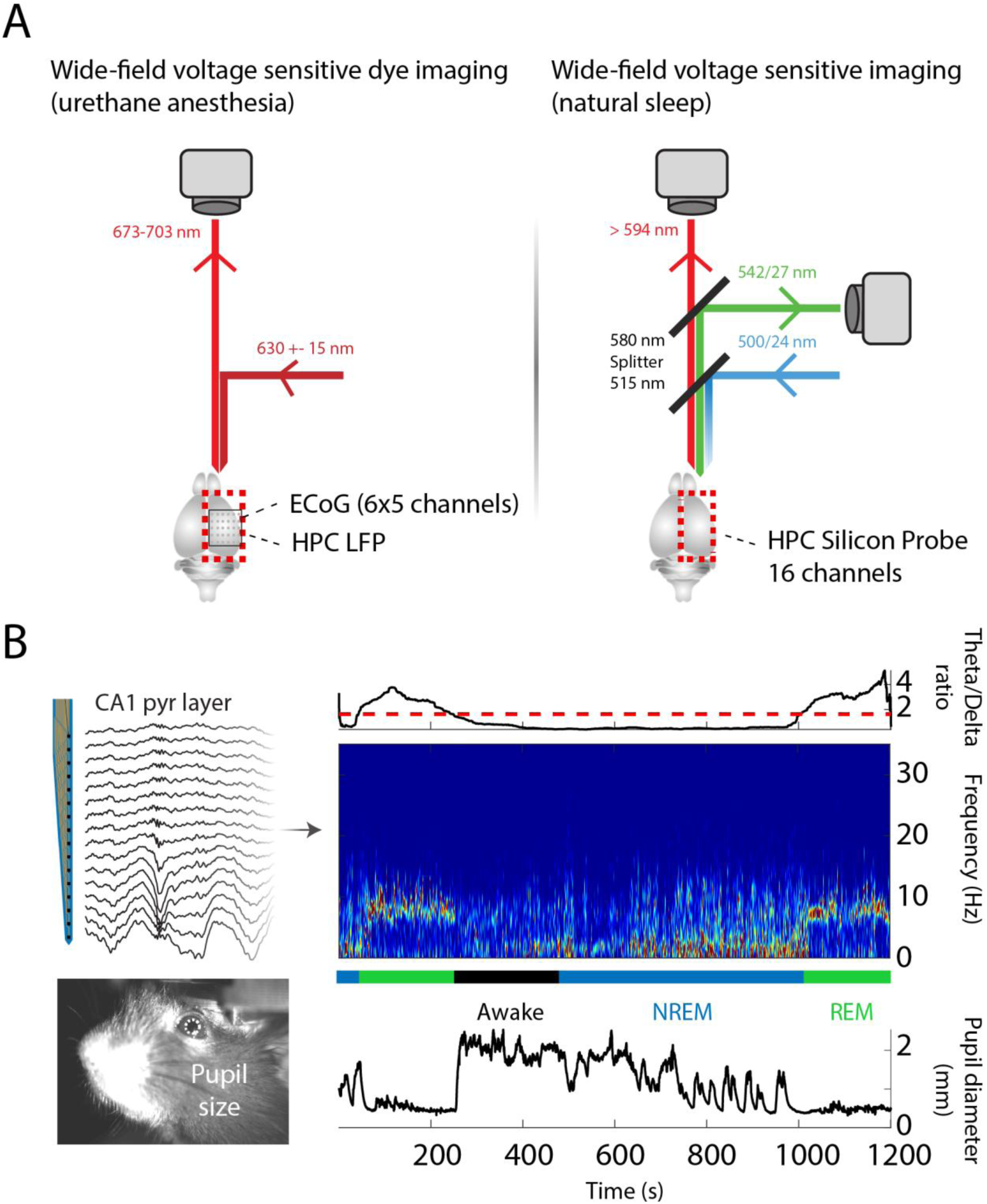
Experimental protocol for simultaneous spontaneous neocortical activity with hippocampal CA1 electrophysiology recordings. **(A)** Schematic of the experimental setups used for the combination of optical imaging with cortical and hippocampal electrophysiology. On the left panel, a CCD camera recorded the fluorescent variability from the superficial neocortical layers. This activity was simultaneously acquired with a transparent ECoG on the superficie of the neocortex and hippocampal CA1 LFP on mice under urethane anesthesia. On the right panel, two PCO edge 4.2 cameras recorded the reflected light from the superficial neocortical layers over the scalp. In combination a high-density silicon probe recorded the hippocampal CA1 profile on mice in natural sleep. **(B)** Example of the sleep state classification for VSI data. Using a channel in the pyramidal layer, we extracted the Theta/Delta ratio to detect the putative periods of awake, NREM and REM sleep. In combination with the facial motion and pupil diameter information extracted using the Deep Lab Cut (DLC), we manually classified the NREM periods which were analyzed in this study.

**Figure S2:**
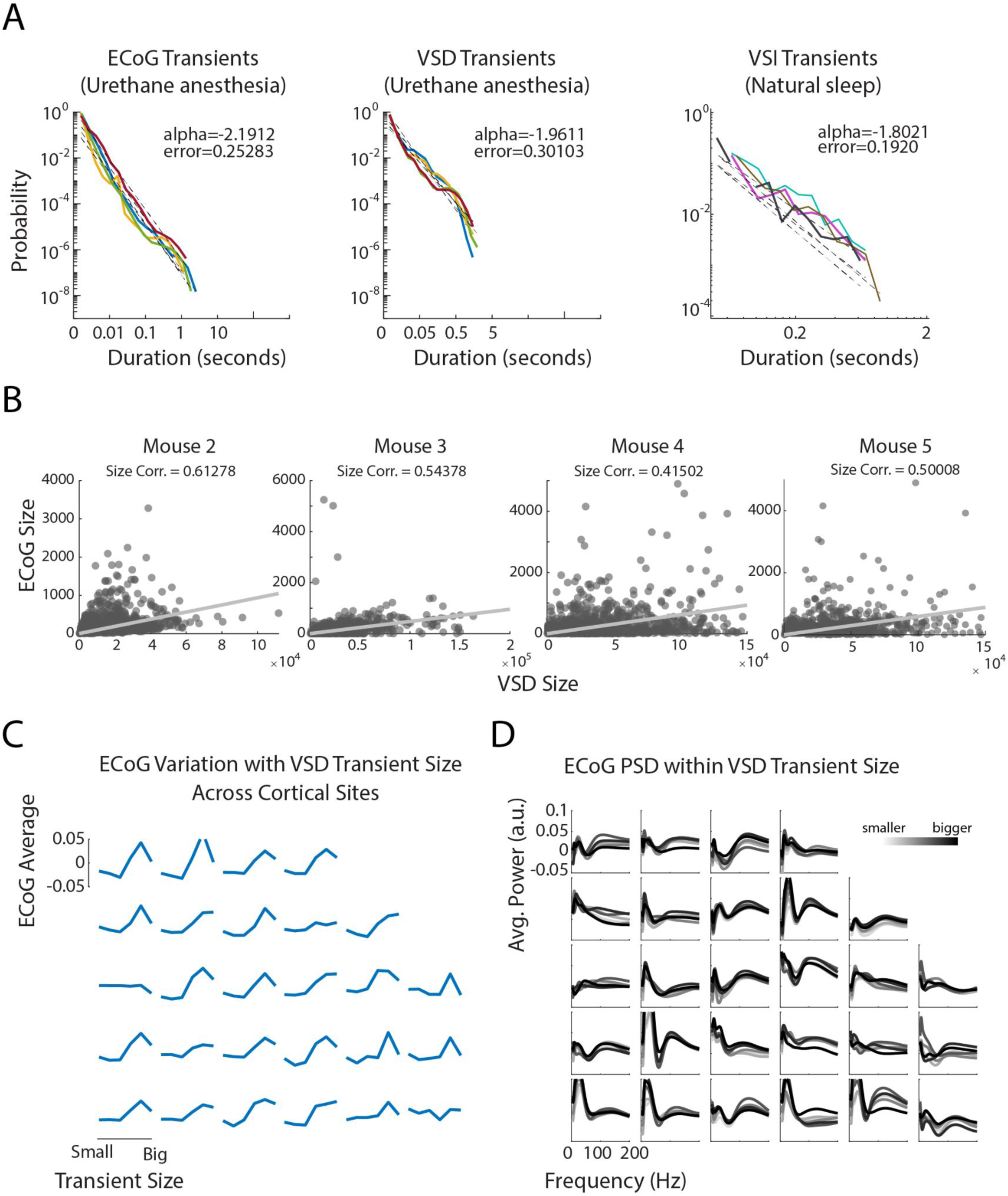
Probability distribution of transient duration in urethane anesthesia and natural sleep. **(A)** Transient probability duration in urethane anesthetized and natural sleep states. The transients’ duration is correspondent from the transient shown in Figure 2. Each color line represents different animals. The black dashed lines represent the linear regression in a logarithm basis of each animal. **(B)** Example of the other animals shown in Figure 2 with the scatter plot of the transient sizes detected by the VSD data and the respective negative of ECoG size. **(C)** Negative power of the ECoG map averaged across animals for different transient sizes detected by the VSD signal. **(D)** ECoG power spectral density during different VSD neocortical transient sizes.

**Figure S3:**
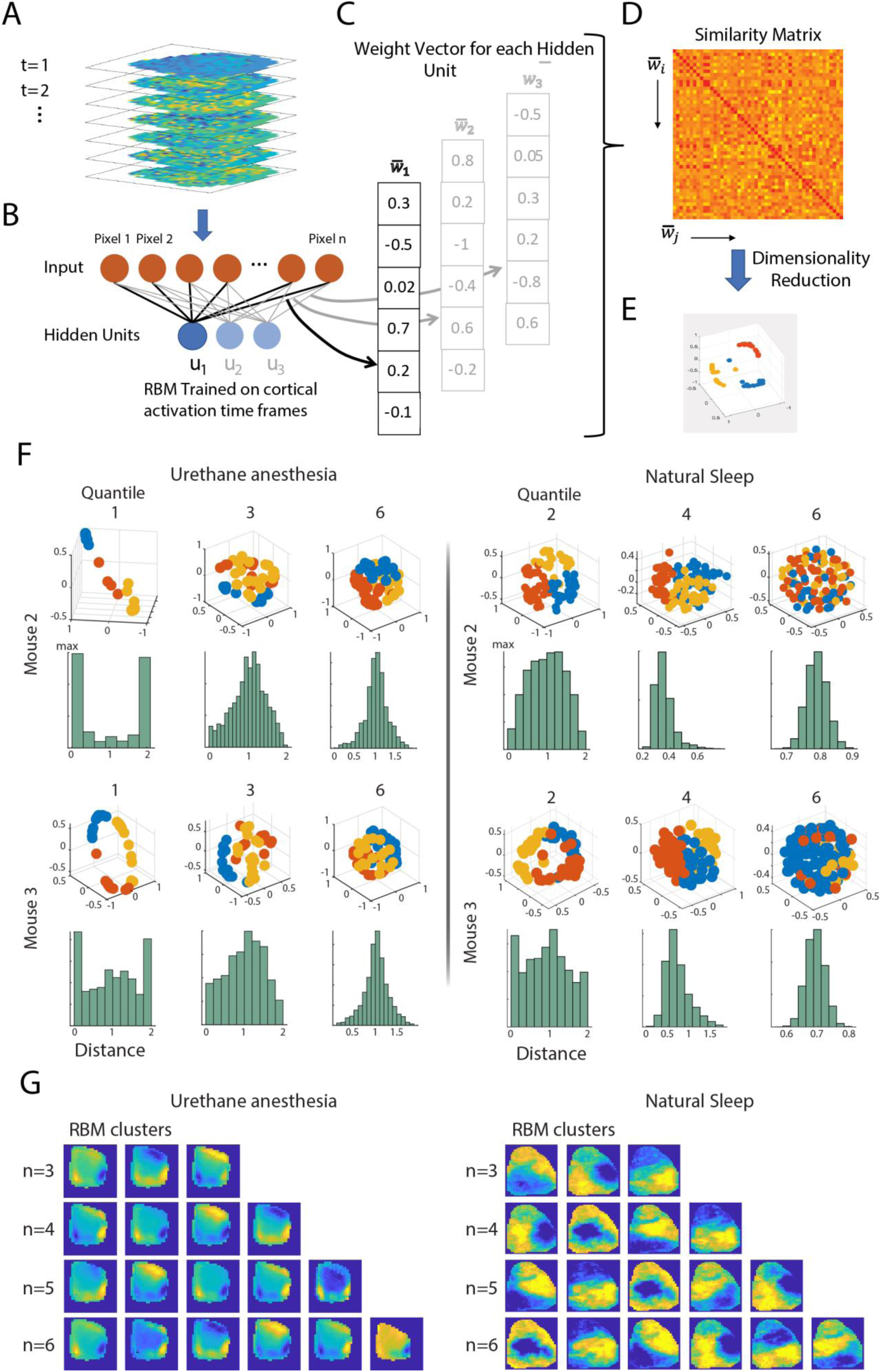
Schematic of the analysis of cortical transients based on a Restricted Boltzmann Machine. **(A)** Single frames from selected temporal intervals are collected and used as the training dataset for the RBM. **(B)** When training the RBM on batches extracted from the input dataset, each frame is presented independently in the input layer. **(C)** At the end of the training procedure, after having achieved convergence, the set of weights associated to each of the hidden units connections is stored separately in a vector. **(D)** A similarity matrix is computed by comparing pairs of hidden units and taking the correlation between their respective weight vectors. This similarity matrix is then used to extract a lower-dimensional representation of the relative similarity of the spatial selectivity (represented by the values of their connection weight) developed by the hidden unit population. **(E)** The same matrix is used to identify clusters of hidden units with similar spatial selectivity. **(F)** Example of 2 mice for the urethane anesthesia and natural sleep datasets showing the RBM weights computed in different quantile groups. The blue, orange and yellow colors represent 3 components separated by k-means classification. The respective green bars show the distributed cosine distance quantile group. **(G)** Spatial distribution of activity corresponding to different k-means component numbers in the first quartile RBM weights.

**Figure S4:**
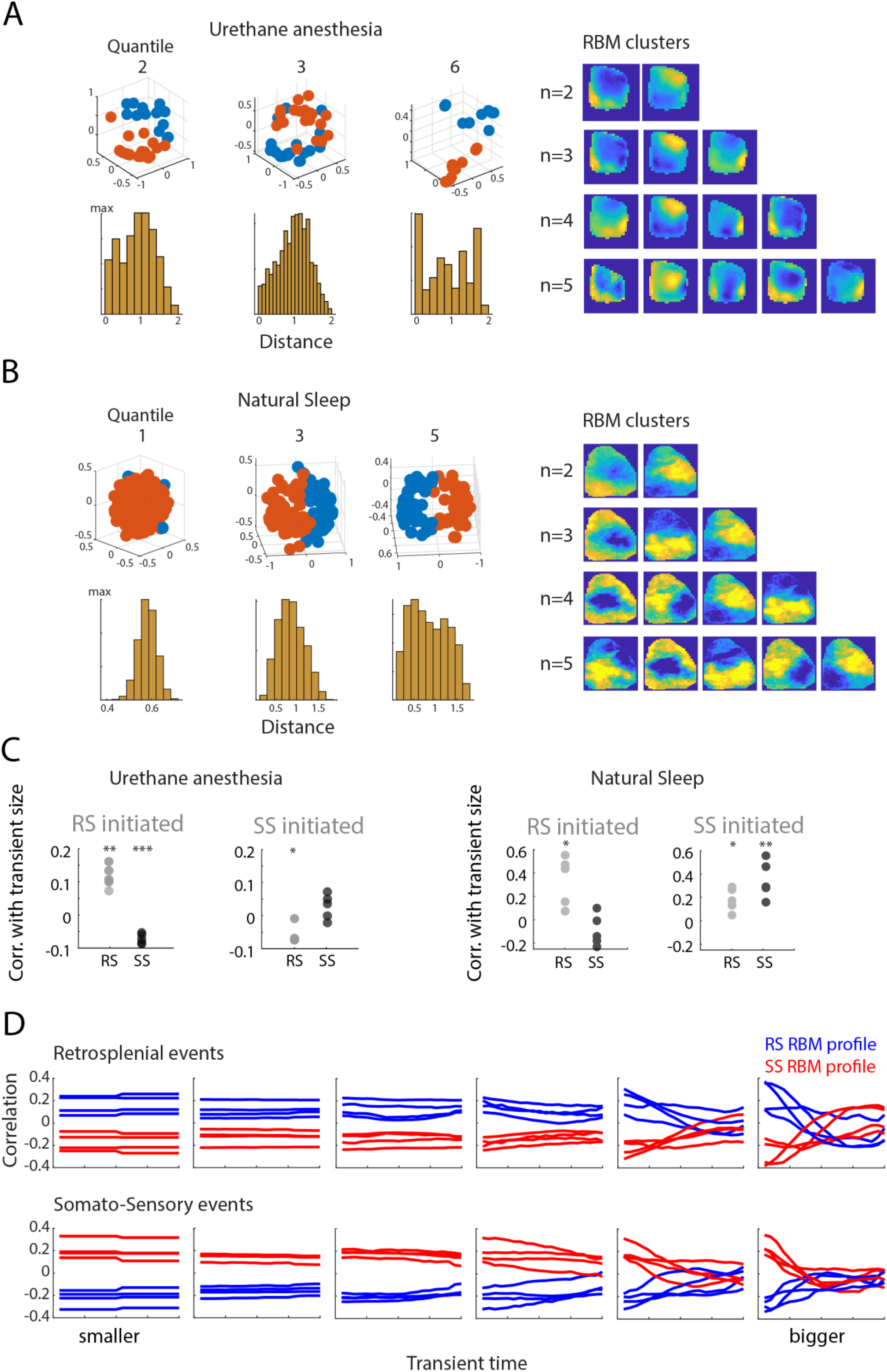
Large transient domains are preceded by Retrosplenial and Somato-Sensory modular activation. **(A)** Example of the RBM weights in a different animal computed on the cortical activation 100 ms before the transient onset in different quantile groups distributed by size for urethane anesthesia data. The respective yellow bars show the distributed cosine distance for each individual pre-transient. **(B)** Same as panel (A) but for an animal in the natural sleep group. **(C)** Correlation mean in RS and SS areas between the transients initiated profile with the RBM extracted networks classified as RS or SS modes (*p<0.05, **p<0.005 and ***p<0.0005; one-sample t-test). **(D)** Correlation of the averaged transients classified as RS (in blue) or SS (in red) with the domain of the transient. Each individual line for each color represents an animal in the urethane anesthesia group (4 lines for RS and 4 lines for SS = 4 animals).

**Figure S5:**
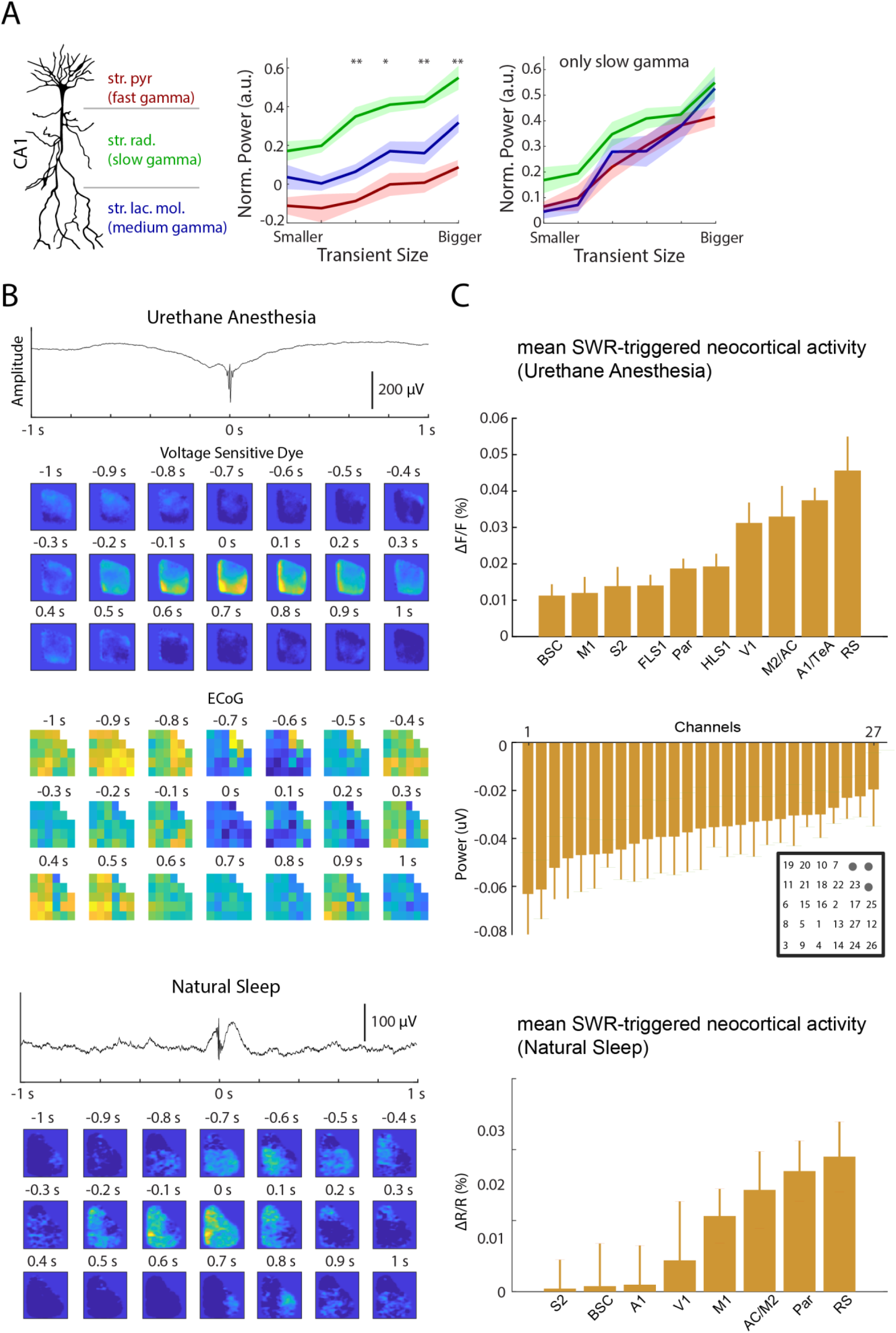
Transient size shows association with Slow Gamma power in stratum radiatum. **(A)** Normalized power for Fast Gamma (100-140 Hz) in s*tratum pyramidale*, Slow Gamma (20-40 Hz) in *stratum radiatum* and Medium Gamma (60-90 Hz) in *stratum lacunosum moleculare* across different transient sizes during natural sleep. A channel in each CA1 layer cited was selected to calculate the normalized power. On the right painel, the same was calculated for specifically Slow Gamma frequency, note that the *stratum radiatum* slightly shows a higher power compared to the other layers. The result points that the influence of the transient sizes correlates mainly with Slow Gamma in the stratum radiatum layer specific. **(B)** Spatiotemporal activity around Sharp-Wave Ripple triggered events in urethane anesthesia (VSD and ECoG) and natural sleep (VSI) states. **(C)** Average of the VSD, ECoG and VSI signals measured from 4×4 or 8×8 pixels (for the VSD and VSI consecutively. ∼0.03 mm2) for the optical imaging activity in each measured area. Bar graphs indicate mean ± SEM.

**Figure S6:**
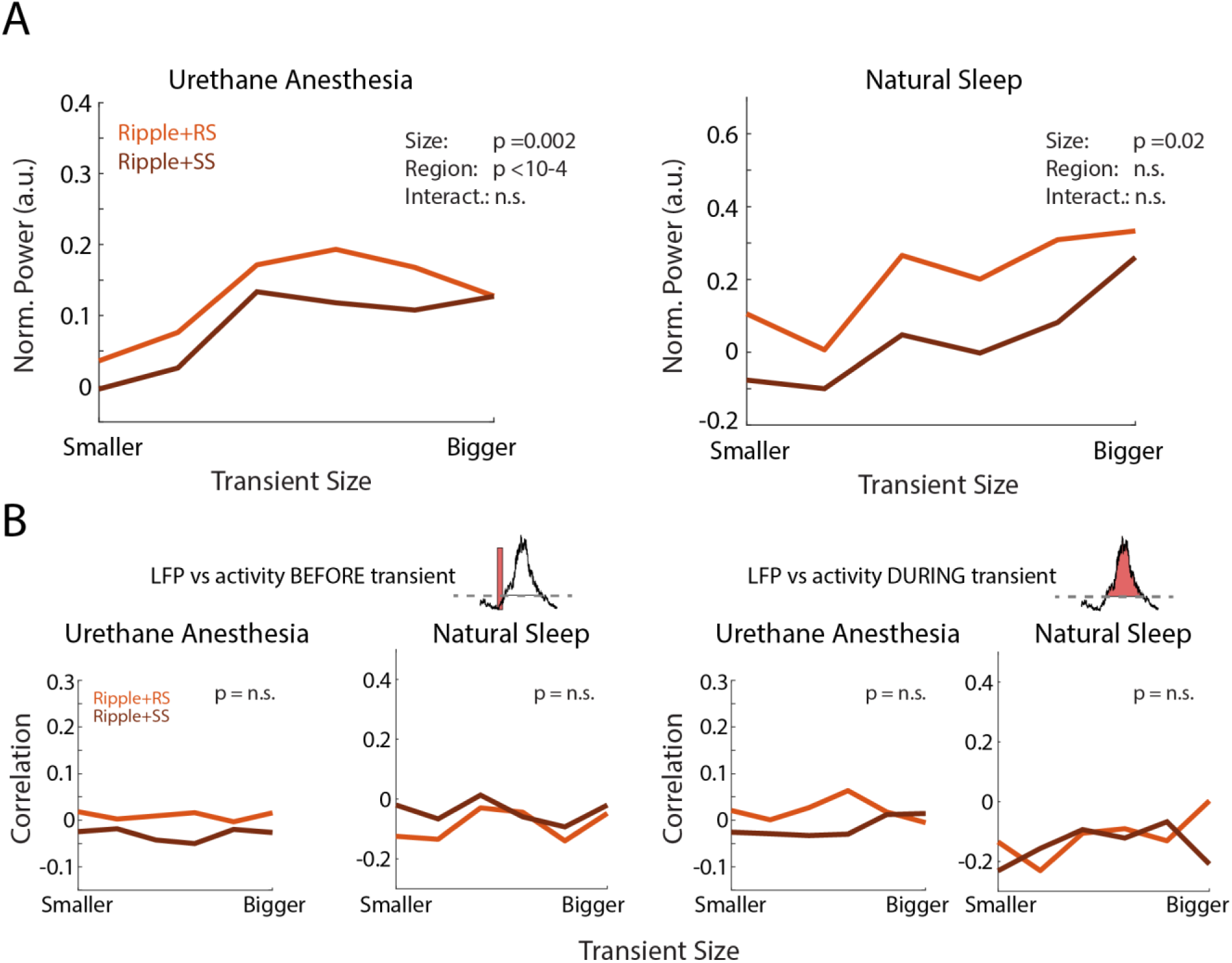
Neocortical transients and Ripple power in the hippocampus. **(A)** Normalized Ripple power (150-250 Hz) for transients triggered by RS and SS across size (two-way ANOVA for ripple in RS vs SS). **(B)** Correlation between Ripple power with RS or SS triggered events within the same block sizes. The left panel is the correlation between the hippocampal CA1 LFP and the activity of the period before RS and SS and the right panel during RS and SS transients.

